# Comparative genomic analysis of a metagenome-assembled genome reveals distinctive symbiotic traits in a Mucoromycotina fine root endophyte arbuscular mycorrhizal fungus

**DOI:** 10.1101/2024.11.27.625642

**Authors:** Joshua Cole, Sébastien Raguideau, Payman Abbaszadeh-Dahaji, Sally Hilton, George Muscatt, Ryan M. Mushinski, R. Henrik Nilsson, Megan H. Ryan, Christopher Quince, Gary D. Bending

## Abstract

**Background:** Recent evidence shows that arbuscular mycorrhizal (AM) symbiosis is established by two distinct fungal groups, with the distinctive ‘fine root endophyte’ morphotype formed by fungi from the sub-phylum Mucoromycotina rather than the sub-phylum Glomeromycotina. While Mucoromycotina AM fungi are globally distributed, there is currently no understanding of the genomic basis for their symbiosis or how this symbiosis compares to that of other mycorrhizal symbionts.

**Results:** We used culture-independent metagenome sequencing to assemble and characterise the metagenome-assembled genome (MAG) of a putative fine root endophyte, which we show belonged to the family Planticonsortiaceae within the order Densosporales. The MAG shares key traits with Glomeromycotina fungi, which indicate obligate biotrophy, including the absence of fatty acid and thiamine biosynthesis pathways, limited enzymatic abilities to degrade plant cell walls, and a high abundance of calcium transporters. In contrast to Glomeromycotina fungi, it exhibits a higher capacity for degradation of microbial cell walls, a complete cellulose degradation pathway, low abundances of copper, nitrate and ammonium transporters, and a complete pathway for vitamin B6 biosynthesis.

**Conclusion:** These differences highlight the potential for contrasting interactions between Mucoromycotina and Glomeromycotina AM fungi with their host plant and the environment which could support niche differentiation and complementary ecological functions.

## Background

Mutualistic symbioses with microbes are exhibited by the overwhelming majority of plant species [1] with mycorrhizas by far the most widespread and well-described association [2]. While mycorrhizal symbioses have evolved multiple times across plant and fungal taxa, they share the common characteristic of the fungus trading nutrients assimilated from the soil, particularly phosphorus and nitrogen, in exchange for carbon from the plant [3]. Additionally, mycorrhizal associations can confer a range of secondary benefits to the plant host, including disease resistance and stress tolerance [4]. Because of these profound impacts on plants, mycorrhizal associations are considered to be major determinants of ecosystem productivity and diversity and to play key roles in directing terrestrial biogeochemical cycling processes. There are four main groups of mycorrhizal associations: arbuscular (AM), ecto-(ECM), ericoid (ERM), and orchid (ORM) mycorrhizas [5], which are distinguished by a number of features including the specific plant and fungal taxa involved, the ecosystems they inhabit, the way in which the fungus colonises and grows within root tissues, and the functional traits of the fungi involved [6].

As a function of time and niche specificity, mycorrhizal symbionts appear to have evolved along a saprotrophy to symbiosis continuum [5, 7]. This directional transition from free-living saprotrophs to obligate symbionts is believed to be driven by the provision of carbohydrates and vitamins from plants in nutrient-poor environments and the subsequent evolutionary success of a symbiotic lifestyle. As a result, key genomic signatures associated with this transition can be identified across this continuum [5]. As fungal symbionts shift away from primarily saprotrophic lifestyles to mutualisms, their genomes typically possess a reduced repertoire of the enzymatic machinery involved in the degradation of soil organic matter (SOM) and plant cell walls [5, 8, 9]. The two youngest mycorrhizal symbioses, ERM and ORM, with origins c 75-90 and 112 mya, respectively [6, 8], involve symbionts with genomic repertoires capable of simultaneously facilitating a saprotrophic and symbiotic lifestyle. However, fungi forming the older AM (origin 450 mya, [6]) and ECM symbioses (origin c 200 mya) possess a much more fine-tuned enzymatic toolkit, noted for a reduced suite of plant cell wall degrading enzymes (PCWDE) along with the presence of specific genes involved in regulating nutrient transport and host plant signalling and metabolism [5, 8].

AM are considered the most common and widespread plant-microbe symbiosis on Earth [10–12], and their presence in fossil roots of the earliest land plants suggests that they were a significant factor in the evolution of land plants. Fungi forming AM have traditionally been placed into the Mucoromycota sub-phylum Glomeromycotina. These fungi are obligate symbionts; as fatty acid auxotrophs they depend on the exchange of plant-derived fatty acids [13, 14]. As a result of long-term host dependence, arbuscular mycorrhizal fungi (AMF) present hallmark traits such as limited ability to break down cellulose, biosynthesise thiamine, or metabolise vitamin B6 derivatives. The genomic inability to express these hallmark traits represent Missing Glomeromycotan Core Genes (MGCGs) [15–17].

Until recently, all AMF were believed to belong to the Glomeromycotina (G-AMF) [12]. It has long been recognised that there are two broad groups of arbuscular mycorrhiza based on morphology, with the ‘coarse endophyte’ G-AMF being the focus of the vast majority of research on the biology and ecology of AMF [18]. Glomeromycotina comprises four orders, the Glomerales, Diversisporales, Paraglomerales, and Archaeosporales, which likely diversified when plants evolved onto land [4, 19, 20]. The second morphological group of ‘fine root endophytes (FRE)’ arbuscular fungi are distinguished from coarse endophytes by their small spores, fan-shaped or palmate entry points, diverging thin hyphae (<2 µm diameter) which in intercellular spaces often form hyphal ropes, as well as intercalary and terminal “vesicles”, approximately 5–10 µm in diameter [18, 21–23]. Interestingly, the arbuscules of FRE appear very similar in structure to those of coarse endophytes [24]. Recent evidence has placed FRE within the sub-phylum Mucoromycotina (M-AMF) [18], indicating that AM symbioses may have evolved independently several times, as is the case for ectomycorrhizal symbioses [25].

To date only one species of M-AMF, Planticonsortium tenue, has been described [23]. However, similarly to G-AMF, root systems are colonised by diverse Mucoromycotina taxa, communities of which are typically comprised of a small number of abundant and common taxa and a high diversity of rare low-abundance taxa [26–28]. Furthermore, distinctive morphologies of FRE have been recognised [21] based on a variety of characteristics, including the roughness, diameter, and branching of hyphae and the size and shape of vesicles. While fungi within the Glomeromycotina are all assumed to be plant symbionts, fungi residing within the Mucoromycotina also include saprotrophs and putative ectomycorrhizal symbionts. Those Mucoromycotina sequences found associated with roots and rhizoids of vascular and non-vascular plants appear to have similar phylogenetic ranges and occur within clades alongside taxa with saprotrophic and putative ectomycorrhizal lifestyles [28, 29].

M-AMF have been detected across multiple continents, spanning a wide range of ecosystems, environmental conditions, and hosts, although in contrast to G-AMF they may be missing from tropical and sub-tropical biomes, and may be favoured by agricultural management practices [22, 27]. While the benefits of G-AMF to the host, particularly for phosphorus nutrition, are well documented, the functional significance of M-AMF within ecosystems is poorly understood [18, 24, 30]. G-AMF and M-AMF typically co-occur and intermingle within root systems [18, 22, 31], but they appear to occupy distinct but overlapping ecological niches [32], with their distribution at the landscape scale differentially determined by temperature, pH, and plant richness [27]. Unlike G-AMF, nutrient availability was found to be an important determinant of M-AMF abundance, which increased with fertility [27]. Furthermore, in contrast to G-AMF, experimental evidence suggests that M-AMF may favour providing the host with access to soil nitrogen, indicating a potential complementary functionality between the two arbuscular symbioses and an explanation for the parallel evolution of symbiotic traits [33, 34].

Comparative genome analysis has emerged as a powerful tool to understand microbial evolution, function, and ecology. However, these approaches largely rely on the availability of cultured microbes or voucher specimens for whole genome sequencing (e.g. [8, 17, 25]). Importantly, culturing microbes can prove time-consuming and problematic as microbial populations have great complexity and many microbes have specific nutritional requirements which cannot be met in culture. This is particularly the case for obligate biotrophs such as symbionts which rely wholly or in-part on the host for their nutritional requirements [35]. As obligate symbionts, G-AMF require a host plant to complete their lifecycle. Despite this, pure cultures of G-AMF have been obtained using root organ cultures, and these have been used to produce inoculum for genome sequencing [36]. The challenges in obtaining and maintaining root cultures have meant that direct genome sequencing of G-AMF cultures has been limited to date. Recently, asymbiotic mycelium and spore production of the G-AMF Rhizophagus clarus has been achieved by supplementing its growth medium with palmitoleic acid, and this could provide a boost to rates of culture-based sequencing [35]. An alternative approach to obtain fungal genomes is sequencing of nuclei extracted from spores [37], although most G-AMF taxa appear to sporulate infrequently, if at all [38].

M-AMF are also putatively obligate symbionts, but because they have only recently been recognised as a distinct group of symbionts, culturing efforts lag behind those of G-AMF. Recently, monoxenic cultures of a Mucoromycotina fungus associated with the clubmoss Lycopodiella inundata have been obtained and shown to form FRE colonisation in a vascular plant [33]. However, while genome sequencing and comparative genomic analysis of a number of saprotrophic and putative ectomycorrhizal Mucoromycotina has been achieved [17, 39], there is no understanding of the genome of M-AMF fungi and how their functional repertoire and mutualistic signatures compare to those of G-AMF, other mycorrhizal symbioses, and saprotrophic and putatively ectomycorrhizal Mucoromycotina. To further complicate the matter, initial comparative analysis of the sister clades indicate that some widely considered hallmark traits of mycorrhizal symbioses may instead represent ancestral markers as opposed to adaptation [17].

Recently there has been great progress in the development of computational tools for the culture-independent assembly of ‘dark’ microbial genomes from metagenome sequencing datasets [40]. Such ‘Metagenome Assembled Genomes’ (MAGs) have been widely used to identify the metabolic capabilities and ecological roles of specific prokaryotes in diverse habitats [41]. MAG construction of eukaryotic microbes is more challenging because of a range of issues including their large sizes, high genome heterozygosity, low abundance in metagenome datasets, and a lack of well-annotated reference genomes for gene annotation [42, 43]. As a result, few studies have constructed eukaryotic MAGs from metagenomes, and these have typically been performed in low-diversity communities (e.g. [42, 44]). A further particular issue with analysis of plant symbionts in situ is their intimate association with plant tissue. This results in co-extraction of host DNA, increasing the sequencing depth required to assemble symbiont genomes.

The aim of the current work was to use culture-independent metagenome sequencing to assemble and characterise the genome of an FRE-forming M-AMF fungus for the first time and compare its genomic repertoire to glomeromycotan fungi and other fungal symbionts. Roots of Trifolium subterraneum were first enriched with FRE using a selective soil sieving technique. Root samples were used for metagenome sequencing, from which a MAG of the FRE was constructed. We compiled a database of 274 fungal genomes across the Ascomycota, Basidiomycota and Mucoromycota, and assigned these to G-AMF, ECM, ERM, OCM, saprotroph, pathogen, parasite, endophyte, lichenic, and mixed lifestyles. Comparative analysis showed that the MAG shares key distinctive characteristics with G-AMF. This includes overlaps in some MCGCs, presence of abundant calcium transporters, and an overall reduced set of plant cell wall degrading enzymes. However, we found a relatively high abundance of microbial cell wall degrading enzymes and evidence of complete cellulose degradation capability, as well as low numbers of nitrogen transporters, suggesting a distinctive niche relative to G-AMF and the other mycorrhizal lifestyles.

## Materials and Methods

### Soil collection and sample preparation

For the metagenome sequencing, plants with the root AMF community enriched in M-AMF were produced following the methods of Orchard et al. (2017a). In brief, soil was collected from the field studied by Orchard et al. (2016) and Orchard et al. (2017a) in the Peel Harvey region in Western Australia, Australia: site coordinates were 32° 48 826’S, 115° 53346’E. Further detail on the location, soil properties, enrichment process and mycorrhizal staining protocol are available in the supplementary information.

### DNA extraction, 18S rRNA gene amplicon sequencing and metagenome sequencing

Twelve samples were selected for 18S rRNA amplicon and metagenome sequencing on the basis of predominance and abundance of M-AMF colonisation as well as quality and quantity of DNA obtained (Table S1). DNA was extracted from root samples using the DNeasy PowerSoilPro Kit (Qiagen) following the manufacturer’s protocol. DNA concentrations were measured using a Qubit 3.0 fluorimeter using the Qubit dsDNA high sensitivity assay kit (Invitrogen) according to the manufacturer’s protocol.

To determine whether the dilution technique had enriched M-AMF, we performed community 18S rRNA amplicon analysis. Glomeromycotina and Mucoromycotina 18S rRNA genes in root samples were amplified using AM-Sal-F and AMDGR primers which amplify a 220-base fragment [28]. PCR reactions, Illumina MiSeq sequencing, and bioinformatic analysis were performed using the approaches outlined in Seeliger et al. (2024).

Long read nanopore amplicon sequencing of the nuclear ribosomal 18S rRNA, 5.8S, and 28S/LSU region of Mucoromycotina was performed on root samples to support construction of the FRE MAG and to determine its phylogenetic relationships. PCR reactions were conducted using AM-Sal-F in combination with RCA95m [45] using the recommended Oxford Nanopore protocol, amplifying a 4.5 kb region. Further detail is available in the supplementary information.

Fast5 files were base called and demultiplexed using the Guppy (2022) basecaller provided by Oxford Nanopore Technologies. The sample was then processed using amplicon_sorter v2022-03-28 [46] which aims to group amplicons at species or genus level (∼95%), output consensus sequences, and quantify the copy number of the representative consensus sequences in the sample. This resulted in 42 consensus sequences, 5 of which were placed into the Mucoromycotina.

To aid detection of M-AMF sequences in the metagenome, a reference 18S rRNA sequence was obtained from DNA extracted from T. subterraneum root samples which were enriched in M-AMF and contained minimal G-AMF. These samples were produced in a similar manner to that described above and are presented in Orchard et al. (2017a) as “Experiment 1”. On average, 90% of the total AMF colonisation in these roots was contributed by M-AMF, presenting the first evidence linking FRE morphology to Mucoromycotina. The AM-Sal-F and RCA95m primers were used to amplify a 4.5 kb region, as described above, which was cloned using the TA cloning kit (ThermoFisher). Partial 18S rRNA analysis using AM-Sal-F and AMDGR primers revealed a clone, FRE4583, whose sequence showed a 100% match both to OTU4 which was associated with FRE morphology in the earlier Orchard et al. (2017a) study and the BEG249 Planticonsortium tenue FRE culture described by Walker et al. (2018) [27]. Furthermore, this and closely related sequences have been detected in multiple studies reporting FRE morphology in plant roots [24, 28]. Primer walking was used to sequence overlapping regions of clone FRE4583 until the full sequence was covered, generating a 4.5 kb fragment.

### Metagenome sequencing and assembly

The twelve root samples were used for Shotgun metagenomic sequencing using the HiSeq 150 paired end protocol to generate between 15 and 47 Gbp sequencing depth. Library construction and sequencing was conducted using Illumina protocols by Novogene UK Ltd.

A collection of reference genomes for T. subterraneum was built from downloading all available sequences from the NCBI at the date of the work (2022). This collection was then used to filter samples DNA using bwa mem v0.7.18-r1243-dirty [47] and samtools v1.10 [48]. To determine FRE depth and breadth of coverage, the filtered metagenome reads were mapped, again using bwa mem and samtools, against the FRE4583 ribosomal operon and the resulting alignment files processed with custom Python scripts. To be sure that a high mean coverage was not an artefact of conserved regions, breadth of coverage was visually inspected. Sample P72 was predicted to have the highest Mucoromycotina abundance, possessing the majority of the operon with a coverage depth of over 100 (Fig. S1); the missing regions are likely due to repeats as we filtered ambiguous mappings. We therefore focused on this sample in our efforts to reconstruct an M-AMF genome, assembling it using the SPAdes assembler v3.14.1 [49] with option -meta. This produced a catalogue of contigs with an N50 of 446 and total length of 1.3 Gbp. We then computed coverage depth profiles for these contigs across all samples by mapping reads with bwa mem, samtools, and bedtools v2.30.0 [50]. This improves the discriminatory power of genome binning through multi-sample coverages whilst keeping the best quality single sample assemblies. The contigs were then binned using these coverage profiles and composition with the binning software CONCOCT v1.1.0 [51] and a minimum contig length of 1000 bp.

The putative M-AMF genome was identified based on a consensus of multiple lines of evidence, see Fig. S2. Initially, we searched for the reference M-AMF ribosomal sequence FRE4583 in our dataset by mapping it to the assembly. A total of 12 unitigs were found with high similarity (>98%) and spanning 75% of the operon without overlap. Since the assembly is fragmented, we decided to explore the assembly graph and more particularly the context of these small unitigs. Out of the 12, only 4 had a context, i.e., they were linked to other unitigs in the assembly graph. The idea behind this approach is that ignoring any miss-assemblies, all genomes have a distinct path in the assembly graph and in particular the M-AMF genome should include a path which passes through those unitigs labelled as ribosomal sequences thus allowing identification of M-AMF. We could not assemble a complete FRE4583 operon in a single contig, which is likely due to strain diversity. Using custom scripts available on GitHub (https://github.com/Sebastien-Raguideau/Graph-tools) we extracted a subgraph encompassing all 10 Kbp around these 4 unitigs, and then visualised all contigs binned by CONCOCT on this subgraph using Bandage v0.8.1 [52], see Fig. S3. In this figure, the 4 unitigs are in red, un-binned unitigs are in gray, and all other colored unitigs are related to different CONCOCT bins. While this figure looks complex, it only corresponds to 0.05% of the whole assembly and refers to a limited radius of 10 kb nucleotides around the ribosomal unitigs, showing a clear proximity of multiple numbered bins.

A total of 12 bins were found in close proximity to M-AMF unitigs. However, on assessing quality, only bin 17 was found to have any marker genes, giving it a completion of 83.7% using the BUSCO v5.2.2 [53] general fungi_odb10 scheme with a size of 37.37 MBp (Table S2). In particular, while contigs from bin 70 were directly co-located with FRE ribosomal sequences, this bin did not possess any marker genes and was distinctly different in terms of coverage from bin 17. It may be that bin 70’s location in the graph is due to a miss-assembly, or it could be that this is due to strain diversity and that bin 70 is strain-specific part of the pan-genome of the FRE which would result in it having a different coverage profile to the core genome represented by bin 17.

To provide additional QC for this putative FRE MAG (bin 17) we used varScan [54] to detect SNVs on all the contigs assigned to it. In Fig. S4 we show the frequencies of these SNVs, and in Fig. S5 their density on contigs. There are a large number of SNVs throughout the genome (Fig. S5) but these are distributed evenly in frequency suggesting that they are likely generated by strain diversity rather than multiple haplotypes. We also computed the distribution of coverage depths of all ORFs predicted by prodigal (Fig. S6). While not sufficient to correctly detect all ORF from a eukaryotic genome, it still suggests that most genes are in a single copy despite the strain diversity.

We used phyloFlash v3.4.2 [55] to quantify abundances of all the SSU rRNAs in the SILVA database [56] and also our 42 target nanopore amplicon 18S rRNA consensus sequences. Filtering any taxa with less than 10 reads mapping across all samples yielded a total of 251 entries including the 42 consensus sequences (Fig. S7). These were normalised by the total sequencing depth of each sample. To compare to these, the coverage depth of bin 17 was calculated for each sample by taking the sum of all nucleotides mapping to all contigs of bin 17 and dividing by the total length of the contigs, followed by normalisation as above. Using Pearson’s correlation, the coverage of bin 17 was compared to the coverage of all 251 taxa detected with phyloFlash. The strongest correlation (r = 0.97 and an adjusted p value of 4.5e-11, Fig. S8) was with one of the FRE consensus sequences (consensus_all_seqs_0_16), thus enabling us to provide finer scale phylogenetic information for bin 17, unambiguously linking it to this M-AMF sequence. The regression coefficient of 63 between coverage profiles (consensus_all_seqs_0_16 and bin 17) provides an indication of the number of ribosomal operons in the genome, a quantity consistent with other fungi [57].

### Reference genome access and ecological lifestyle annotation

To compare the MAG to known fungi, a database of reference fungal genomes was constructed from JGI MycoCosm (Table S3). The fungal species, representing the three phyla—Ascomycota, Basidiomycota, and Mucoromycota—were chosen based on two criteria. First, fungal genomes which had been included in previous comparative genomics analyses of mycorrhizal fungi [8, 25] were included since the ecology of these fungi had already been curated. Secondly, the database was supplemented with Mucoromycota genomes available in MycoCosm (https://mycocosm.jgi.doe.gov/mucoromycota/mucoromycota.info.html) in May 2023. The associated predicted fungal proteomes were accessed and downloaded in May 2023. Fungal genomes were categorised according to their ecological lifestyle (Table S3). We assigned 146 genomes as saprotrophic, 42 as ECM, 29 as pathogenic, 23 as mixed lifestyle, 12 as G-AMF, 9 as endophytic, 5 as parasitic, 4 as ERM, 3 as OCM, and 1 as lichenic. Within this total, we included five Mucoromycotina genomes. Full descriptions of the ecological assignment process and the Mucoromycotina genomes are available in the supplementary materials.

### Phylogenetic assessment

Phylogenetic assessment of FRE4583 and those consensus sequences from root samples which were assigned to the order Endogonales (including all_seqs_0_16) was carried out through 18S rRNA gene alignment. These sequences were added to the Endogonomycetes 18S rRNA alignment from Tedersoo et al. (2024) using MAFFT v7.525 [58]. The resulting alignment was manually checked and refined in MEGA11 v11.0.13 [59]. The best fitting model for our alignment was selected using IQ-TREE v2.3.6 and the option “-MFP” [60, 61]. A phylogenetic tree was generated using the model “TIM2+F+R5” with 1000 approximate likelihood ratio tests and 1000 ultrafast bootstraps [62, 61]. The tree was subsequently visualised in FigTree v1.4.4 [63].

In addition to the 18S rRNA alignment, fungal phylogeny was inferred from genome sequences using a concatenated protein marker alignment. A set of 434 protein markers identified by the JGI as pan-fungal markers for inferring high-level fungal phylogeny was used to compare the phylogenetic placement of the MAG and all the reference fungal genomes within our curated database. To identify these marker proteins, an HMM set was downloaded from https://github.com/1KFG/Phylogenomics_HMMs/tree/master/HMM/JGI_1086.

Phylogenomic markers were extracted from fungal proteomes using the above HMM, and a subsequent phylogenetic tree was built using PHYling v0.9.0 (available at: https://github.com/stajichlab/PHYling) under default settings. The tree was then visualised in R using ggtree v3.6.2 [64].

### Annotation of MAG and reference genomes

Our comparative analyses focused on Carbohydrate-Active enZymes (CAZymes), lipases, proteases, transporters, and small-secreted proteins. Full details on the methodology used to identify these groups within the genomes is available in the supplementary information.

### Comparative genomics analyses

All comparative genomics analyses were conducted using R v4.2.2 [65], and all visualisations were generated with ggplot2 v3.4.0 [66]. Functional annotation of EC numbers was assigned using DeepECtransformer v1.0 [67] and the ECRECer web platform [68].

## Results

### Description of FRE in the roots

The appearance of the FRE in the roots of the twelve samples chosen for metagenomic sequencing was consistent with the description of the genus *Planticonsortium* by Walker et al. (2018) into which they placed *Glomus tenue* (previously generically referred to in the literature as fine endophyte or fine root endophyte [17] (see photographs in: Fig. 1 herein; Figure 2 and Figure S1 in Orchard et al., 2017a and Thippayarugs et al. (1999)). In particular, entry points were characterised by structures with “feather-like, fan-shaped, palmate or multilobed with finger-like projections” [22] which then led to fine, diverging hyphae in cortex cells and, often, plentiful finely-branched arbuscules. Small intercalary vesicles and terminal vesicles were also common. Droplets of lipids were often evident within hyphae and arbuscules. In some instance, colonisation in the cortex was too dense to clearly distinguish the structure of individual arbuscules.

**Figure 1.**
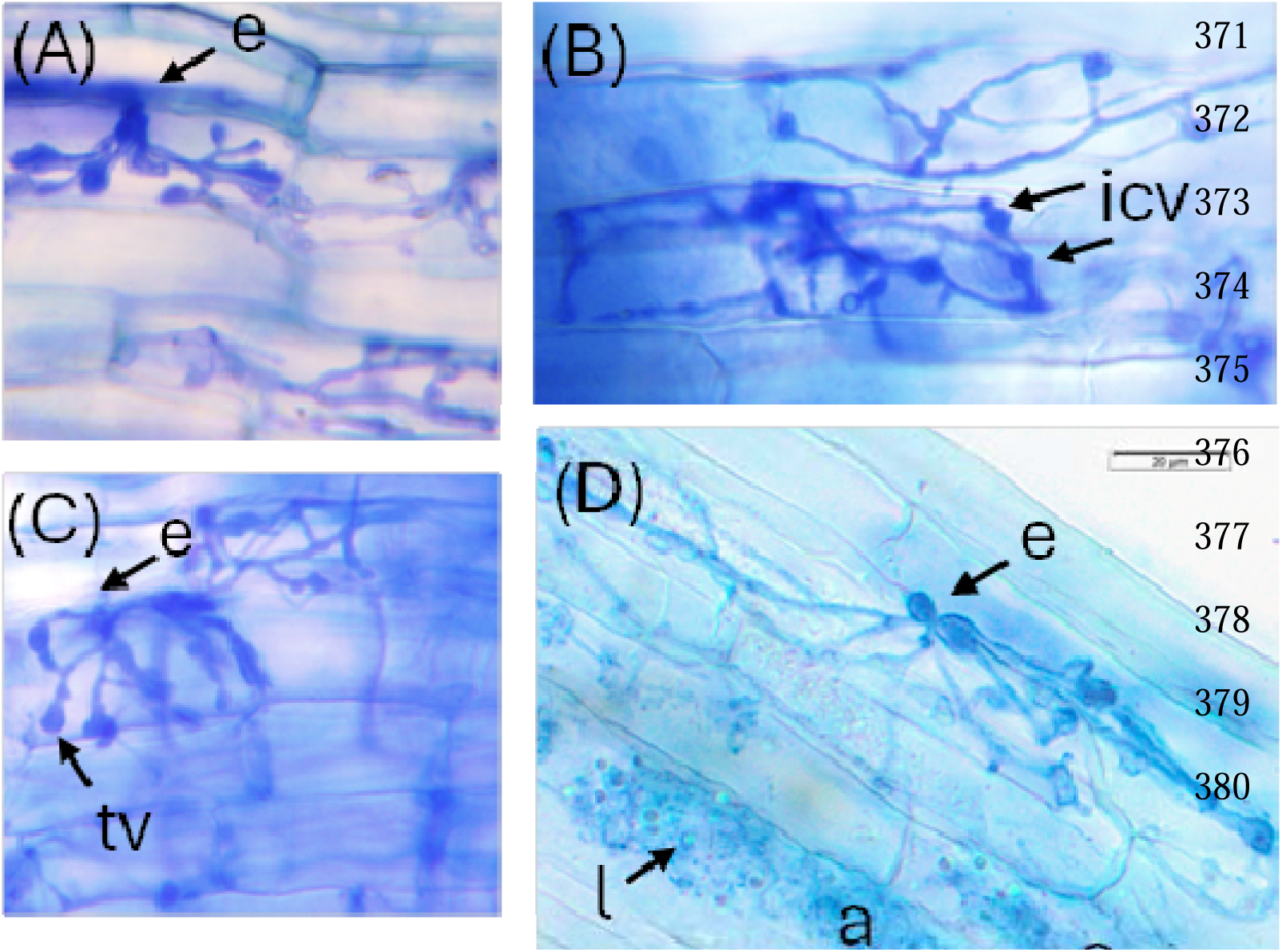
Colonisation by arbuscular mycorrhizal fungi (AMF) with fine root endophyte (FRE) morphology in the roots of Trifolium subterraneum from: (A, B, C) the experiment from which roots were chosen for metagenomic sequencing and (D) roots enriched in FRE which were used as the reference 18S rRNA sequence (FRE 4583) of Mucoromycotina-AMF (Orchard et al. (2017); experiment 1). Evident are the characteristic thin hyphae of FRE morphology, as well as entry points (e), several cells filled by an arbuscule (a) which also contained lipid droplets (l), terminal vesicles (tv) and intercalary vesicles (icv). (Photo credit: Payman Abbaszadeh-Dahaji A, B, C; Suzanne Orchard D)

In the samples used for amplicon and metagenome sequencing, the percentage of total root length colonised by AM ranged between 26.8 and 72.5% for AMF with FRE morphology and 0 to 16.3% for G-AMF, producing a proportion of AMF colonisation by FRE between 62.2 and 100%. 18S rRNA Miseq sequencing of Glomeromycotina and Mucoromycotina communities in root samples revealed that the proportion of Endogonales amplicons ranged between 25 and 82% (Table S1).

### Phylogeny and Features of Fungal Genomes

In this study, we constructed a putative FRE genome, referred to as the ‘MAG’. This MAG was identified as a putative M-AMF through two lines of evidence. First, the contig bin was colocated on the assembly graph with unitigs that we could assign to the 18S rRNA gene of a known M-AMF reference (FRE4583). Secondly, its abundance correlated across the 12 samples in the study with an 18S rRNA sequence ‘consensus_all_seqs_0_16’, which we showed to be very closely related to FRE4583 (Fig. 2). The bin itself did not contain a complete 18S rRNA sequence. This is unsurprising as this gene is well known for failing to assemble, largely due to it being comprised of strongly conserved regions that repeat throughout the genome, failing to assemble into substantial contigs. The MAG measured 37.37 Mbp with a genome coverage varying between 1x and 568 depending on the sample. The N50 was found to be 4928 with a GC content of 41.1%. Finally, its completion was assessed to be 83.7% with only 1.5% contamination (Table S2). We identified a total of 10,872 proteins. Of these proteins, we were able to assign 3,358 proteins with a functional annotation, comprising 218 CAZymes, 121 lipases, 280 proteases, 13 small secreted proteins, and 2,726 transporters.

While we acknowledge that the revised phylogeny of the Mucoromycotina subphylum produced in Tedersoo et al. (2024) lacks type specimens and is grounded in part on dark taxa, as a poorly captured branch of the fungal tree of life, we deem this an appropriate basis to apply the phylogeny of our MAG. 18S rRNA analysis placed consensus_all_seqs_0_16, and by proxy the MAG, within the order Densosporales of the Mucoromycotina, corresponding with the Densosporaceae sensu Desirò et al. (2017) (Fig. 2). This order is associated with FRE morphology [22, 39]. Further to this, strong support values allow us to confidently place the MAG in the newly proposed family Planticonsortiaceae [69]. Within this family, the MAG was placed with very high similarity to the known M-AMF reference sequence, FRE4583 (Fig. 2). The MAG was closely related to three other Mucoromycotina 18S rRNA sequences obtained from the root samples (consensus_all_seqs 0105, 0122, and 0260) and to environmental sequences obtained from the thalli of early diverging plant sequences, e.g., MH174585 and MH174471. The wider Planticonsortiaceae clade included further sequences from the thalli of non-vascular plants, sequences not annotated beyond the phylum level associated with soil from across Europe (EUK sequences), and one sequence associated with the rhizosphere of Populus tremuloides (EF23703), a known arbuscular mycorrhizal host [70]. Phylogenetic analysis therefore placed the MAG apart from other members of the Mucoromycotina with putative ecology from the order Endogonales, including the putative ectomycorrhizal Mucoromycotina member Jimgerdemannia spp. [29] and the saprotrophic Endogone sp. and Vinositunica sp. which have a putative saprotrophic or ectomycorrhizal lifestyle [71].

**Figure 2:**
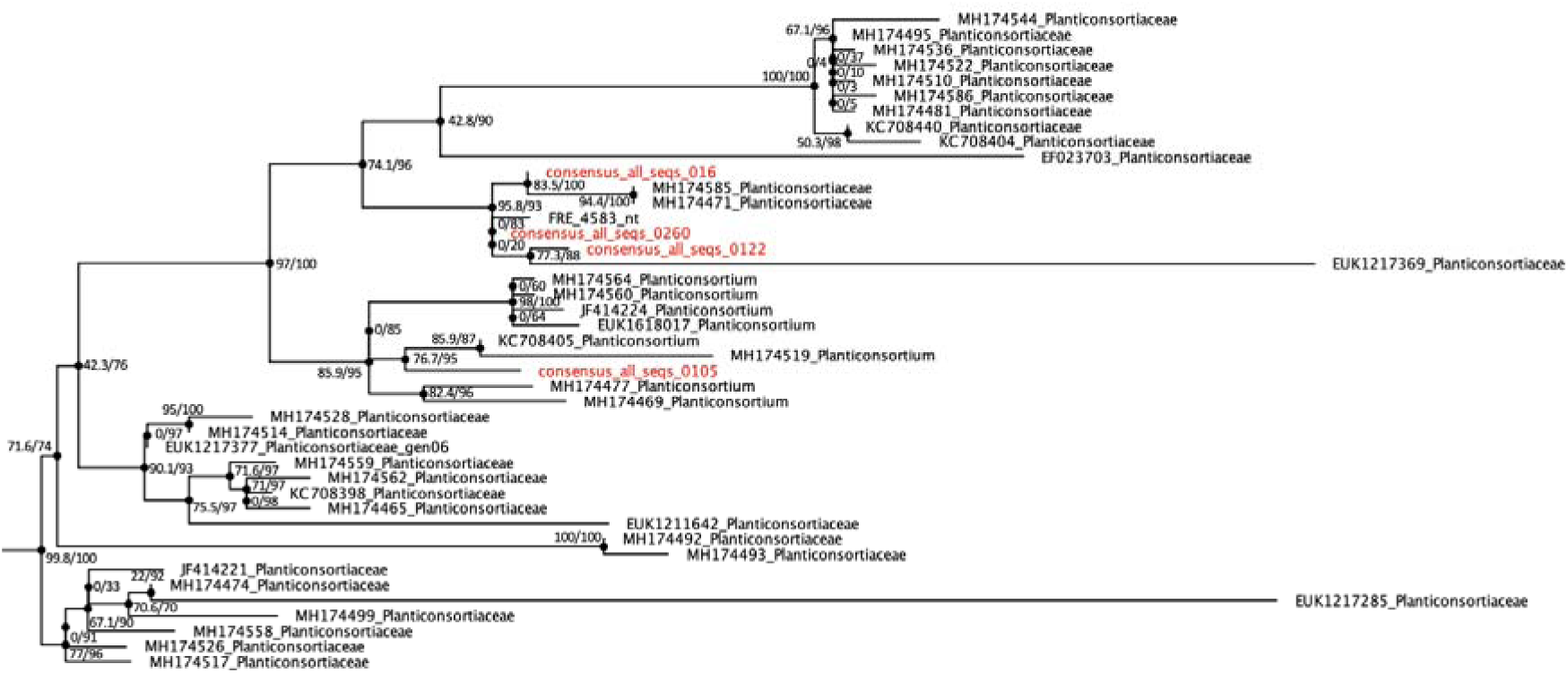
Phylogenetic assessment of the Mucoromycotina-AMF reference sequence FRE 4583 and ‘consensus_all_seqs_0_16’, an 18S rRNA gene sequence associated with the MAG, and related 18S rRNA sequences amplified from root samples (consensus_all_seqs_0122, 0105 and 0122, highlighted red), in the context of the Tedersoo et al. (2024) Endogonomycetes SSU alignment, showing the family Planticonsortiaceae. The phylogenetic tree was generated using IQ-TREE, the model “TIM2+F+R5”, 1000 ultrafast bootstrap replicates, and approximate Likelihood Ratio Tests. Branch support is presented as a percentage (aLTR support/bootstrap support).

Similarly, the protein marker alignment placed the MAG within the Mucoromycota (Fig. 3). In our phylogenetic analysis, the MAG forms a sister group to all other Mucoromycotina species except for Calcarisporiella. This placement suggests that the MAG represents an early-diverging lineage within the Mucoromycotina. Due to the lack of available genomes from the order Densosporales, we cannot definitively assign the MAG to this order, but its basal position indicates it may be related to early diverging Mucoromycotina lineages, possibly including the Densosporales.

**Figure 3:**
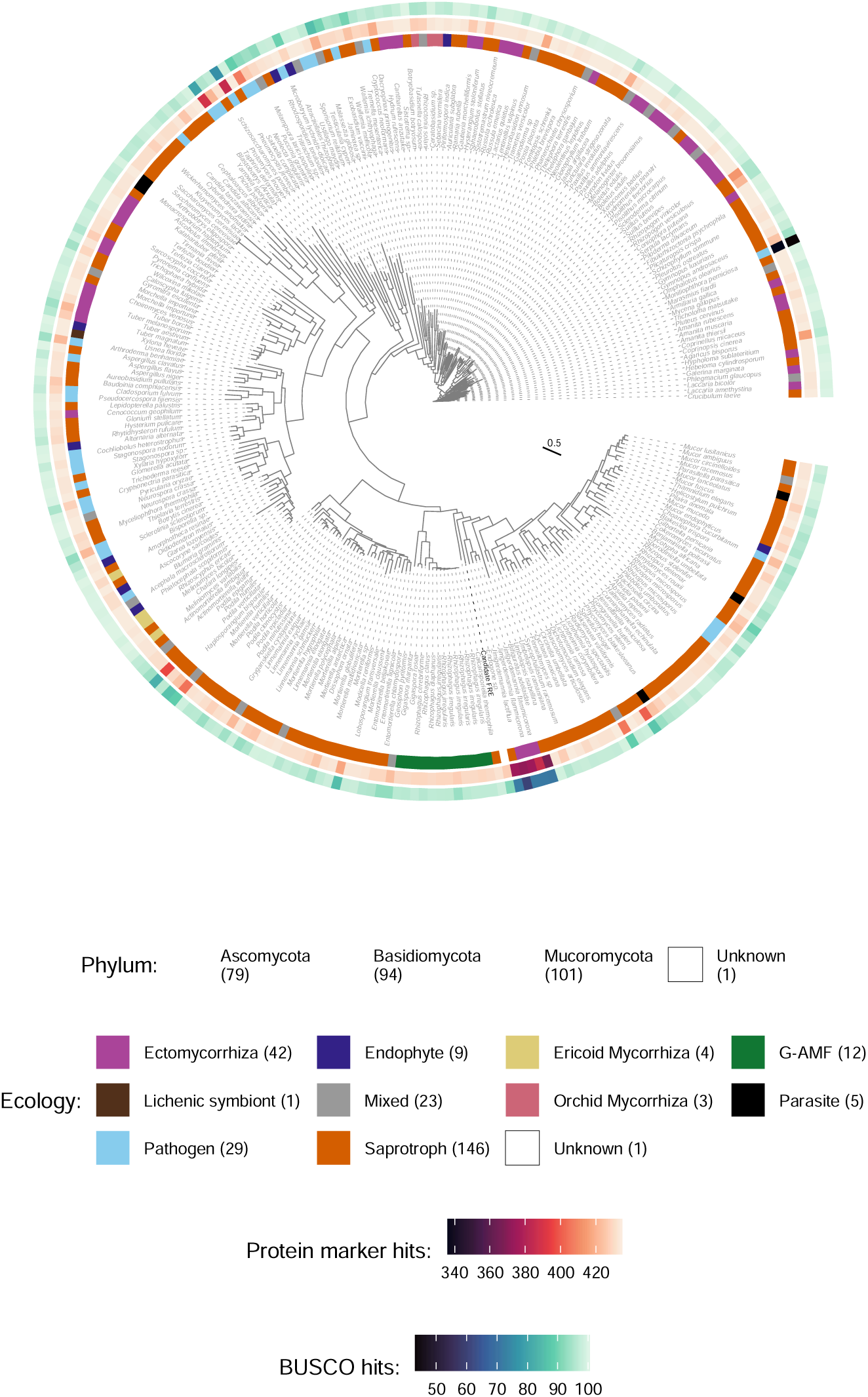
Phylogenetic placement of the MAG (shown here as Candidate FRE) based upon a set of 434 protein markers identified by the JGI as pan-fungal markers for inferring high-level fungal phylogeny. Associated ecological group, protein marker hits and BUSCO hits are shown.

### Genomic Profile of the MAG

To contextualize the gene content of the MAG, we compared the CAZyme, protease, lipase, and transporter profiles to genomes within the reference dataset that represented the saprotrophy-symbiont continuum. Due to the MAG’s early-diverging position within the Mucoromycotina (Fig. 3), and the absence of available genomes from some lineages such as the Densosporales, we highlighted the inclusion of five available genomes from the Endogonales for comparative purposes (Fig. 4-8). While the 18S rRNA and genome-wide analyses indicate that the MAG is a sister group to other Mucoromycotina species (excluding Calcarisporiella), the Endogonales genomes provide the closest available genomic context within this subphylum. This comparison allows us to explore potential functional attributes and evolutionary relationships within the Mucoromycotina. The ecological lifestyle had a significant effect on genome content across all four categories (p < 0.01) (Fig. 4). For the CAZymes, the ecological lifestyles fell into two groups—G-AMF and ECM, ERM and OCM —with SAP bridging the two (Fig. 4a). The MAG contained a low number of CAZymes (218), which was similar to the mean of the ECM (240), but slightly higher than that of the G-AMF (175). A similar number of CAZymes were found in the MAG compared to Endogonales genomes (149 to 208). The ERM genomes contained by far the largest median number of CAZymes across the lifestyles (629). Only the ERM possessed a significantly (p < 0.01) larger number of lipases than the other ecological lifestyles (Fig. 4b). The MAG possessed 121 lipases, lower than the means of the G-AMF (189), ECM (238), and SAP (263), OCM (268). Similarly, the Endogonales genomes all contained low numbers of lipases (72 to 132). The ERM genomes had significantly (p < 0.01) higher mean numbers of proteases per genome (457) than the other lifestyles (Fig. 4c). The MAG had similar numbers of proteases (280) compared to G-AMF (305), ECM (286), SAP (271), and to the Endogonales genomes (229 to 298). The G-AMF and ERM lifestyles had significantly higher (p < 0.05) numbers of transporters per genome relative to the other genomes. The MAG (2726) had a substantially lower number of transporters compared to G-AMF (3900); however, it contained more than the Endogonales genomes (1932 to 2388) that sat in the lower end of the dataset (Fig. 4d). Across the dataset, the saprotrophs contained an especially high degree of intra-group variability in numbers of all four profiles compared to the other ecological lifestyles (Fig. 4). The number of CAZymes, lipases, proteases, and transporters in the MAG were also comparable to the endophyte, parasite pathogen, and mixed lifestyles (Fig. S9). The MAG contained 13 small secreted proteins, a middling amount for the dataset (Fig. S9).

**Figure 4:**
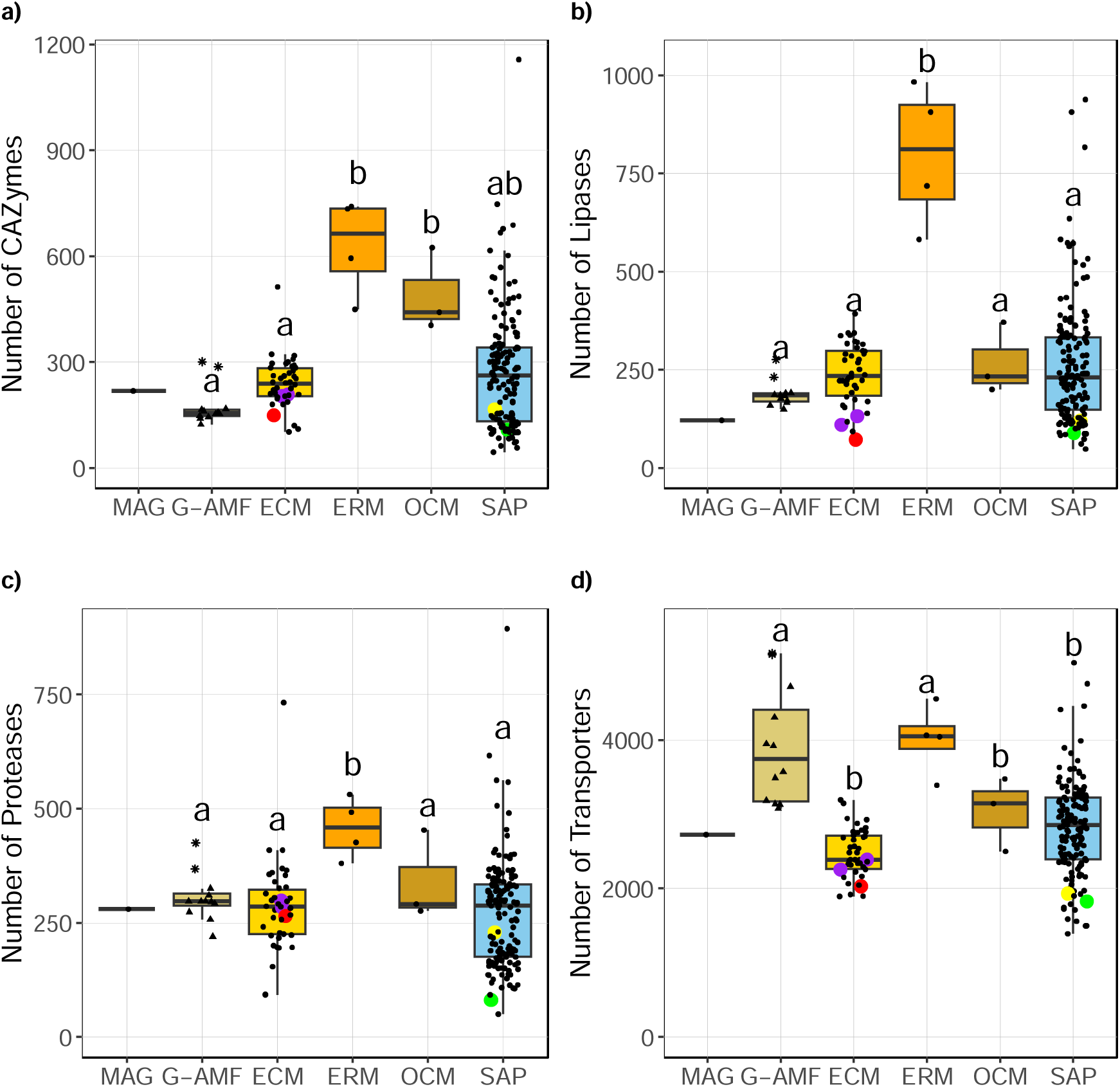
Number of CAZyme, lipase, protease, and transporter predicted proteins per genome distributed by ecological lifestyle. Lifestyle is colour coded. Coloured data points represent selected Mucoromycotina species for comparison: yellow for ‘Endsp1’, red for ‘Jimlac1’, green for ‘Bifad1’, and purple for both ‘Jimfl_AD_1’ and ‘Jimfl_GMNB39_1’. Data point shape within the G-AMF distinguishes between the Diversisporales (star) and Glomerales (triangle). Different letters indicate significant differences between lifestyles as determined by a Kruskal-Wallis test (p < 0.01) followed by a post-hoc Dunn’s test (p < 0.05).

### CAZyme Profile

CAZyme proteins associated with the degradation of plant and microbial cell walls were identified to investigate the saprotrophic repertoire of the MAG, in the context of the mycorrhizal saprotrophy-symbiont continuum. The ERM (252) and OCM (215) genomes possessed a significantly higher (p < 0.01) mean number of PCWDEs than the G-AMF (32), ECM (59), and SAP (77) genomes (Fig. 5a). The MAG contained a similar number of PCWDE (37) as the G-AMF genomes and the genomes of the Endogonales (33 to 44). ERM genomes contained a significantly (p < 0.01) higher mean number of MCWDE (109) than the saprotrophs and other mycorrhizal lifestyles (24 to 59) (Fig. 5b). The G-AMF genomes had a significantly (p < 0.05) lower number of MWCDE (24) compared to the mean ECM (45), OCM (59), and SAP (48) genomes. The MAG had 53 MCWDE, higher than the median number of all other lifestyles except the ERM (Fig. 5b, Fig. S10). A similar number of MCWDE was detected across the Endogonales genomes (36 to 53).

**Figure 5:**
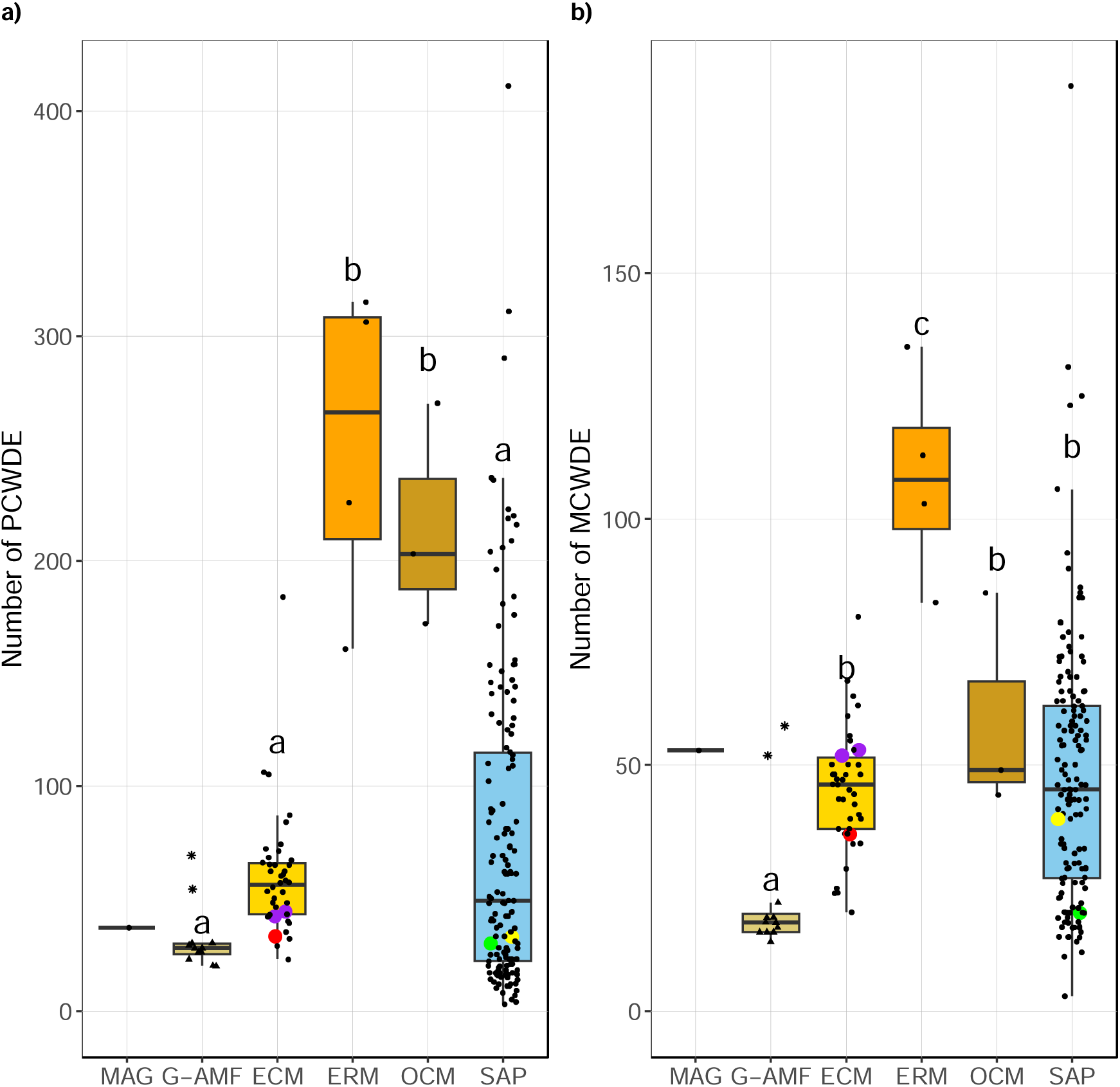
Number of plant cell wall degrading enzyme (PCWDE) and microbial cell wall degrading enzyme (MCWDE) predicted proteins per genome distributed by ecological lifestyle. Lifestyle is colour coded. Coloured data points represent selected Mucoromycotina species for comparison: yellow for ‘Endsp1’, red for ‘Jimlac1’, green for ‘Bifad1’, and purple for both ‘Jimfl_AD_1’ and ‘Jimfl_GMNB39_1’. Data point shapes within the G-AMF distinguish between the Diversisporales (star) and Glomerales (triangle). Different letters indicate significant differences between lifestyles as determined by a Kruskal-Wallis test (p < 0.01) followed by a post-hoc Dunn’s test (p < 0.05).

Patterns of PCWDE distribution across the mycorrhizal associations were similar (Fig. 6). ERM and OCM genomes had significantly (p < 0.01) higher mean numbers of cellulases, hemicellulases, and pectinases than the G-AMF and ECM genomes. Relative to the mean G-AMF, the MAG had notably higher numbers of cellulases (18 compared to 8), but similar numbers of hemicellulases (11 compared to 7) and pectinases (4 compared 2). The Endogonales genomes had similar numbers of cellulases (11 to 13), hemicellulases (10 to 16), and pectinases (4 to 12) as the MAG. The numbers of lignin degrading enzymes were significantly higher in the ERM (31, p < 0.01) relative to the other mycorrhizal lifestyles, which all possessed similar numbers: ECM (13), G-AMF (15), OCM (9), and SAP (12). Notably the MAG had a very low number of lignin degrading enzymes (4) compared to all other lifestyles, this trend was consistent across the Endogonales genomes (2 to 8).

**Figure 6:**
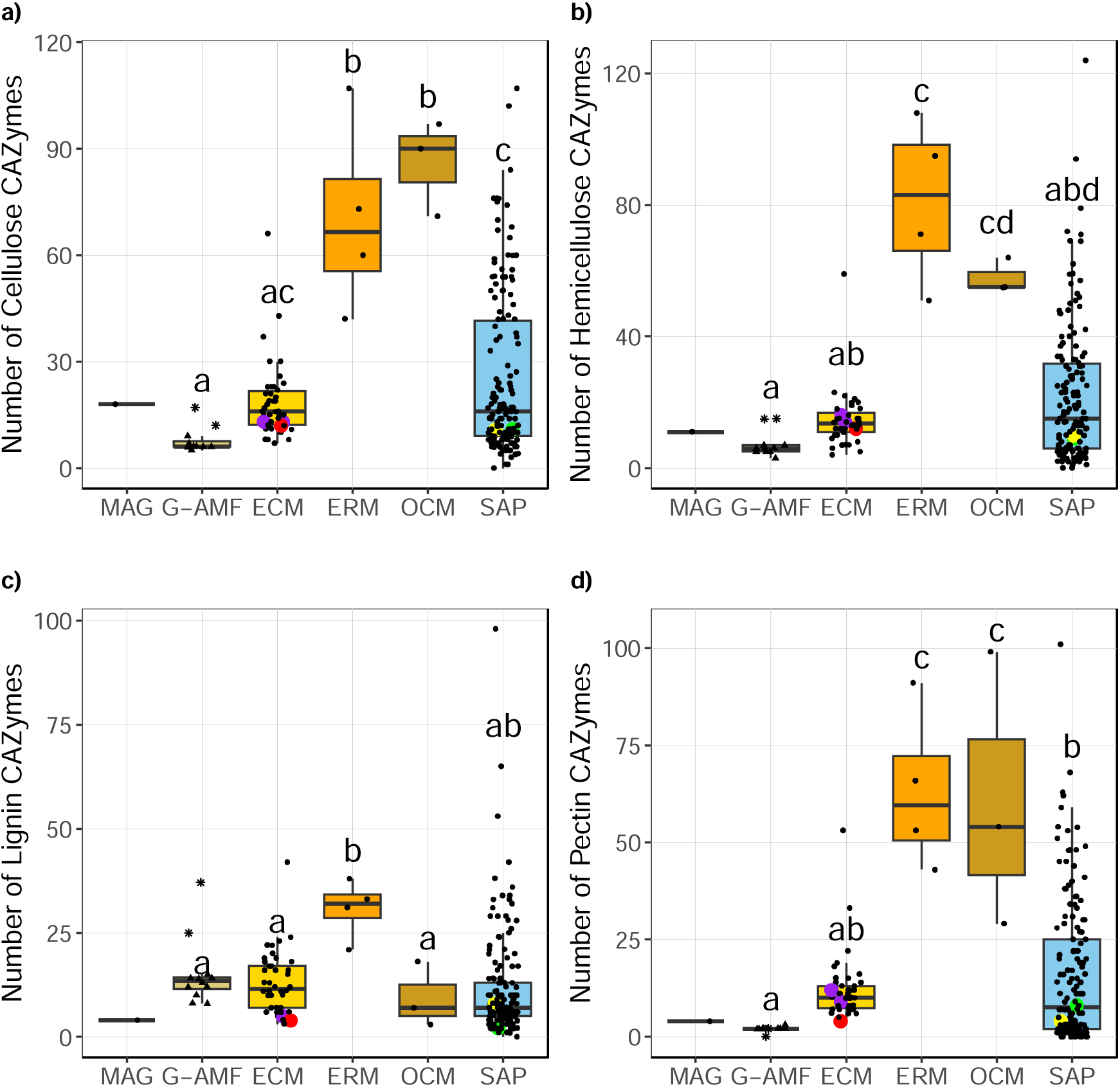
Number of cellulose, hemicellulose, lignin, and pectin degrading predicted proteins per genome distributed by ecological lifestyle. Lifestyle is colour coded. Coloured data points represent selected Mucoromycotina species for comparison: yellow for ‘Endsp1’, red for ‘Jimlac1’, green for ‘Bifad1’, and purple for both ‘Jimfl_AD_1’ and ‘Jimfl_GMNB39_1’. Data point shapes within the G-AMF distinguish between the Diversisporales (star) and Glomerales (triangle). Different letters indicate significant differences between lifestyles as determined by a Kruskal-Wallis test (p < 0.01) followed by a post-hoc Dunn’s test (p < 0.05).

Cellulases in the MAG genome comprised 8 families, with 6 copies of AA9 being the most abundant (Table S4). The Endogonales genomes contained an average of 6 cellulase families, with GH9 being the most abundant in all four genomes, of which the MAG contained 3. However, not a single copy of AA9 was identified within the Endogonales genomes (Table S4). The G-AMF genomes only contained an average of 3 families, with the Glomerales containing a mean of just 6 cellulases. However, the Diversisporales contained a much higher number of cellulases (17 and 12), including 7 and 6 of AA9 and 6 and 3 copies of AA3, respectively (Table S4). The MAG contained a full suite of cellulose degradation enzymes: Endoglucanase, β-glucosidase, and exoglucanase (Table S5). However, with the absence of β-glucosidases in all G-AMF genomes and only two containing a single copy of exoglucanase, none of the G-AMF genomes possessed the full cellulose degradation pathway (Table S5). Hemicellulases in the MAG comprised the majority of the remaining PCWDEs, with 7 families and 5 copies of GH31 being the most abundant (Table S4). Similarly, the Endogonales genomes contained an average 8 families, with GH31 being the most abundant. The G-AMF genomes contained an average 3 families, with GH31 and GH5_7 being the most abundant (Table S4). No copies of GH29 or GH95 were found within the G-AMF genomes. Similarly, no copies of GH95 were found within the MAG. However, a single copy of GH29 was present (Table S4).

Pectinases in the MAG were limited, with only 4 copies across 3 families: 2 GH35s, 1 GH2, and 1 PL1_7 (Table S4). The Endogonales genomes contained an average of 4 families, with the most abundant being GH28 and GH35 (Table S4). Within the G-AMF genomes, no evidence for pectinases was found within ‘Gigmar1’. Apart from that, all other genomes contained only 2 or 3 copies of GH35 (Table S4).

The MAG contained 3 families of lignin degrading enzyme: 2 AA1, 1 AA1_2, and 1 AA2 (Table S4). This trend was conserved across the Endogonales genomes, in which lignin degrading enzymes comprised of 3 families, with AA1 being slightly more abundant (Table S4). The G-AMF genomes, while only comprising an average 3 families, contained a much larger number of lignin degrading enzymes. The Diversisporales contained 37 and 25 lignin degrading enzymes, including 35 and 23 copies of AA1 respectively. The Glomerales contained an average of 12 lignin degrading enzymes per genome, with an average of 9 counts being attributed to AA1 (Table S4).

The G-AMF genomes had a smaller mean chitinase repertoire (16) than ERM (35, p < 0.01), OCM (25, p < 0.05), and SAP (26, p < 0.01) (Fig. 7a). However, the MAG had one of the highest numbers of chitinases across the genome database (38), roughly double the mean number found in both the G-AMF and ECM (21). However, there was considerable variation in the numbers of chitinases across the G-AMF genomes, with the two Diversisporales genomes containing 38 and 29 counts respectively, more comparable to the MAG. The Endogonales genomes similarly contained high mean numbers of chitinases (29 to 42) compared to other genomes in the dataset. Over half of the MAG’s chitinase repertoire was attributed to just two families, CE4 and GH18, with 16 and 13 copies respectively, a trend also seen across the Endogonales genomes (Table S4). The G-AMF genomes largely comprised of CE4 chitinases, however, the Diversisporales genomes also contained a large number of GH18 (13 and 10), very few of which (0 to 2) were found in Glomerales (Table S4). The MAG contained a fully comprehensive set of chitin degrading enzymes (Table S5). Similarly, all but one of the G-AMF genomes contained a full chitin degradation suite (Table S5).

**Figure 7:**
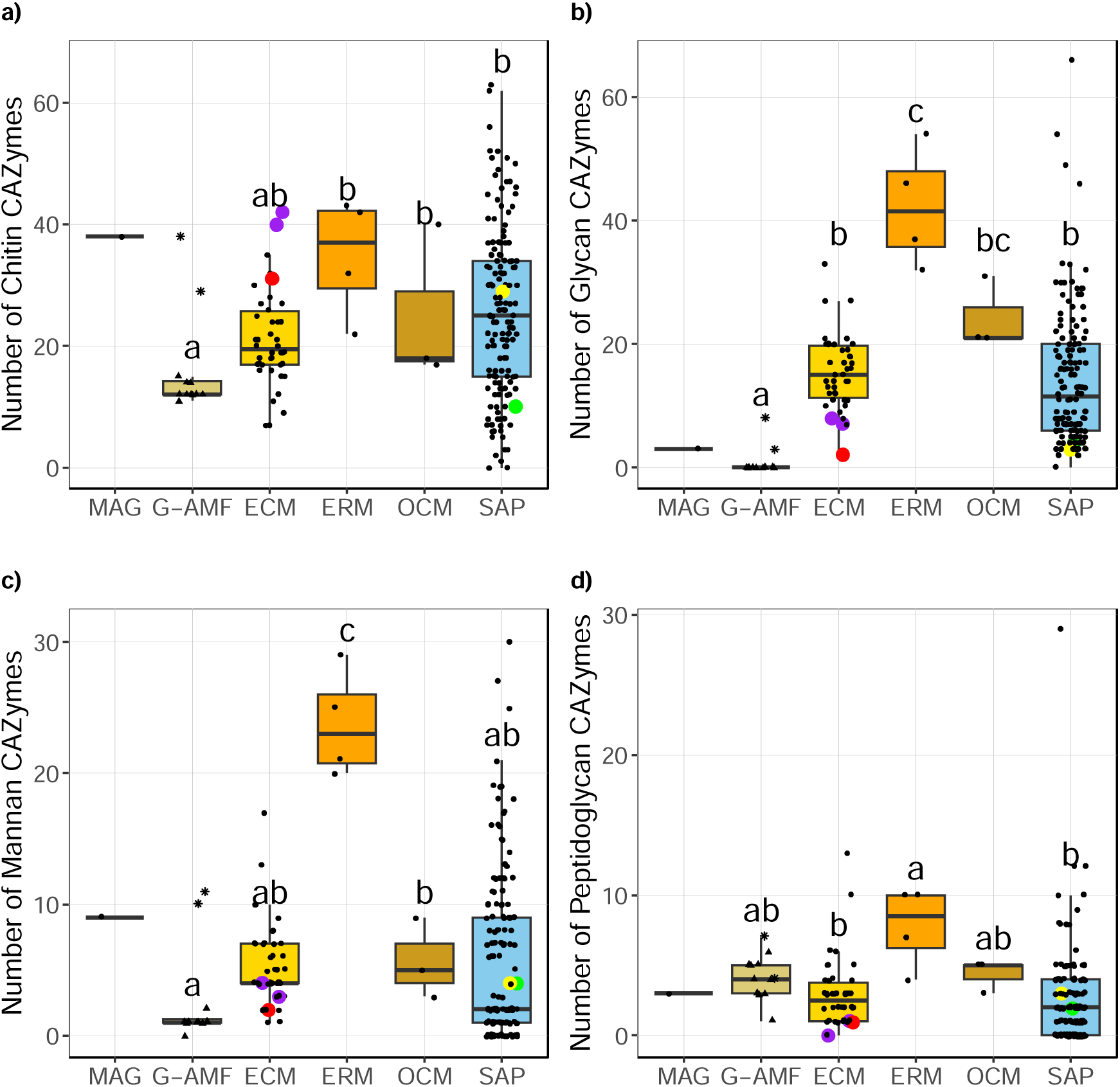
Number of chitin, glycan, mannan, and peptidoglycan degrading predicted proteins per genome distributed by ecological lifestyle. Lifestyle is colour coded. Coloured data points represent selected Mucoromycotina species for comparison: yellow for ‘Endsp1’, red for ‘Jimlac1’, green for ‘Bifad1’, and purple for both ‘Jimfl_AD_1’ and ‘Jimfl_GMNB39_1’. Data point shapes within the G-AMF distinguish between the Diversisporales (star) and Glomerales (triangle). Different letters indicate significant differences between lifestyles as determined by a Kruskal-Wallis test (p = 0.012 for chitin, p = 0.024 for peptidoglycan, and p < 0.01 for the remainder) followed by a post-hoc Dunn’s test (p < 0.05).

ERM genomes contained a higher mean number of mannanases (24) than the saprotrophic and other mycorrhizal lifestyles (3 to 6) (Fig. 7c). However, the MAG contained a substantially higher mean number of mannanases (9) than the G-AMF (3) and ECM (5), and also the Endogonales genomes (2 to 4). Similarly, there was a range of mannanases across the G-AMF genomes with the two Diversisporales genomes containing 11 and 10 counts respectively, more comparable to the MAG (Table S4). The MAG again contained a comparatively large set of genes for mannan degradation, with 5 copies of GH92, 3 copies of GH125, and 1 copy of GH76. However, this represents the one substrate-specific group that was not as conserved across the chosen Endogonales genomes. None of the Endogonales genomes contained copies of GH76, while all contained 1 or 2 copies of CE17 (Table S4). Additionally, while three of the four contained GH92, it was only in single-copy instances. The G-AMF genomes showed a range of mannanases within their genomes, and particularly the Diversisporales genomes alone contained many copies of GH92. All but one Glomerales genome only contained 1 or 2 copies of GH125, with no mannanase identified in the ‘RhiirC2’ genome (Table S4).

The MAG contained no genome-specific cell wall degrading enzymes families. However, a poorly described GH151 family, “α-L-fucosidase enzyme encoding protein”, was only identified within 3 of the Endogonales genomes (Jimfl_AD_1, Jimfl_GMNB39_1, and Endsp1) and a far-removed saprotrophic basidiomycete, Dacsp1 (Table S4). The MAG possessed similar numbers of glycanases and peptidoglycanases to G-AMF and Endogonales genomes (Fig 7b, d and supplementary information).

### Transporters

Since the ability to transport nutrients and assimilate with the host plant is core to the lifestyle of a fungal symbiont, we compared substrate-specific transporters between the ecological lifestyles. Only the ERM genomes contained a statistically higher (p = 0.044) number of phosphate transporters than the other genomes (Fig. 8a). The MAG contained a relatively low number of phosphate transporters (14), although this was not too dissimilar from the mean of the G-AMF (18) and the same as the mean ECM genome (14). Similarly, the Endogonales genomes possessed low numbers of phosphate transporters (13 to 15) (Fig. 8a). The G-AMF genomes had a significantly higher (p < 0.01) number of calcium transporters (55) compared to the other mycorrhizal and saprotrophic lifestyles (Fig. 8b, which ranged between 26 and 39), and also the endophyte, parasite, pathogen, and mixed lifestyles (Fig. S13). The MAG showed a similar number of calcium transporters (50) to the G-AMF, while the number of calcium transporters in the Endogonales genomes varied considerably (25 to 47) (Fig. 8b). There was no significant difference (p = 0.48) in the number of copper transporters across the mycorrhizal and saprotrophic genomes, with the mean varying between 13 and 17 (Fig. 8c). However, the MAG contained only 6 copper transporters, one of the lowest of all genomes (Fig. 8c, Fig. S13). Of the Endogonales genomes, ‘Jimlac1’ had the second smallest number of copper transporters (7) while the rest ranged from 8 to 13 (Fig. 8c). The G-AMF had a significantly higher (p < 0.01) number of nitrate transporters (7) compared to the other mycorrhizal and saprotrophic lifestyles (Fig. 8d). The MAG contained a low number (2) of nitrate transporters relative to the mean G-AMF, although this was the same as the mean ECM (2) and similar to the ERM (3), OCM (1), and SAP (1). There was considerable variation in the mean number of nitrate transporters within the Endogonales genomes (0-5) (Fig. 8d). The ERM genomes had a significantly higher (p < 0.01) mean number of ammonium transporters (11) than all but the G-AMF genomes (9) (Fig. 8e). The MAG contained 5 ammonium transporters, slightly higher than the median ECM (4) and OCM (4). The MAG contained an especially low number of glycine transporters (1), the same as the average ECM (1) and OCM (1) genome (Fig. 8f). The ERM contained the highest mean number of glycine transporters (5), whereas the G-AMF (3) and SAP (4) were not significantly different to any of the other ecological lifestyles (Fig. 8f).

**Figure 8:**
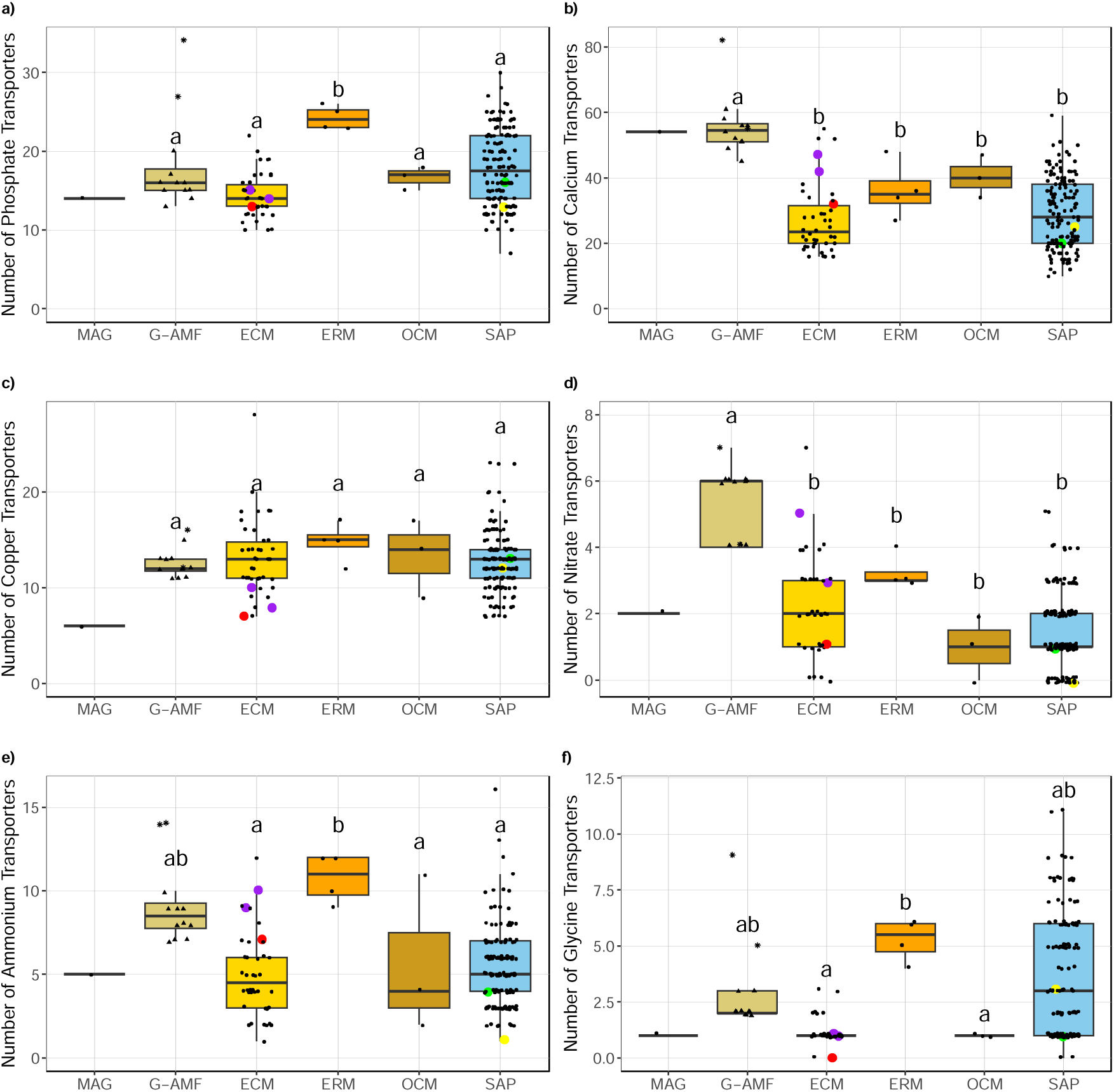
Number of phosphate, calcium, copper, nitrate, ammonium, and glycine transporters per genome distributed by ecological lifestyle. Lifestyle is colour coded. Coloured data points represent selected Mucoromycotina species for comparison: yellow for ‘Endsp1’, red for ‘Jimlac1’, green for ‘Bifad1’, and purple for both ‘Jimfl_AD_1’ and ‘Jimfl_GMNB39_1’. Data point shapes within the G-AMF distinguish between the Diversisporales (star) and Glomerales (triangle). Different letters indicate significant differences between lifestyles as determined by a Kruskal-Wallis test (p = 0.028 for phosphate, p = 0.62 for copper, p = 0.041 for nitrate, and p < 0.01 for the remainder) followed by a post-hoc Dunn’s test (p < 0.05).

### Hallmark Missing Glomeromycotan Core Genes

We analysed both the MAG and G-AMF genomes for enzymes linked to pathways that are missing from typical G-AMFs, often referred to as ‘Missing Glomeromycotan Core Genes’ (MGCG). As fatty acid auxotrophs, G-AMF have been shown to lack the presence of cytosolic fatty acid synthases required for fatty acid biosynthesis. We found no evidence of proteins directly associated with the fatty acid synthase required to facilitate fatty acid biosynthesis within either the G-AMF genomes or the MAG. We did identify 3-oxoacyl-ACP reductase in all genomes, including 10 within the MAG. (Table S5). Additionally, neither the MAG nor the G-AMF genomes possessed the ability to biosynthesis thiamine, a hallmark of G-AMF genomes. Specifically, we found no evidence of the presence of thiamine phosphate synthase or hydroxymethylpyrimidine kinase (Table S5).

The ability to synthesis all derivatives of the vitamin B6 pathway was present within the MAG. This included a copy of pyridoxine kinase (2.7.1.35), the most prolific enzyme within the vitamin B6 metabolism pathway. Only one enzyme, pyridoxine 4-dehydrogenase, was missing from the pathway, while an indirect three-step metabolism for the conversion of pyridoxal to pyridoxine was present. Conversely, no G-AMF genome possessed a full vitamin B6 metabolism pathway, with no evidence of pyridoxine kinase being found across the group (Table S5).

## Discussion

In the current work, we assembled the genome of a putative FRE-forming AMF from a root metagenome. Methodologically, this is a relatively unique achievement: fungal MAGs have been obtained previously from host-associated environments and niche environments, but not from complex environments, such as the rhizosphere [72–74], or from similar fungal symbionts. To achieve this, we had to augment standard binning methods with assembly graph visualisations and coverage correlations to known M-AMF 18S rRNA genes. We believe that this method has many advantages over isolation and culturing, in particular, it is less time-consuming and in principle may be more reflective of the actual community present, particularly if multiple species are present.

We acknowledge that our assembled genome size of 37.37 Mb may underestimate the actual genome size. Short reads can, at times, have inherent difficulties in resolving repetitive regions and complex structures, especially when the repeats exceed the length of the reads. Consequently, repetitive sequences or duplications may collapse, presenting a smaller genome [75]. The early diverging G-AMF Paraglomus occultum genome exhibited a very similar sized (39.6 Mb) genome with very small numbers of repeats, highlighting that a large genome is not a prerequisite of an AM lifestyle [76, 77]. Furthermore, many Mucoromycotina genomes of varying ecological lifestyles are represented by genomes of similar size and limited complexity [78]. There is conflicting evidence as to whether arbuscular mycorrhizal genomes are diploid or simply ‘diploid-looking’ examples of heterokaryosis [79]. Since short-read assemblers cannot phase chromosomes, in the event that M-AMF is a diploid organism, while the genome contents will be assembled correctly, it will be as a haploid chromosome.

Phylogenetic analysis of the correlative 18S rRNA marker, consensus_all_seqs_0_16, showed that the MAG sits within the order Densosporales of the Mucoromycotina. The MAG represents the first published genome of both the order Densosporales and the family Planticonsortiaceae. Due to the absence of sequenced genomes from the Densosporales—the order to which the MAG likely belongs—the only available genomes from the Mucoromycotina for comparison are those of putative ectomycorrhizal and saprotrophic fungi from the order Endogonales. Although these genomes provide valuable comparative insights, the MAG is phylogenetically distinct from these taxa, and a reasonable degree of dissimilarity is evident. The MAG was closely related to environmental sequences from the rhizoids of liverworts, which form arbuscular-like symbioses involving Glomeromycota and Mucoromycotina [80]. To date, understanding of the phylogeny of fungi which form mycorrhizas with FRE morphology in vascular plants has been restricted to short 18S rRNA fragments. Similarly, these studies have confirmed close phylogenetic relatedness with sequences from liverwort thalli [22, 26–28], suggesting these fungi form associations with both vascular and non-vascular plants, with a role in N and P nutrition of the plant host [22, 33]. Mucoromycotina sequences associated with plant roots may span both the Densosporales [18, 29, 39] and Endogonales [28], and furthermore diverse assemblages of fungi from these orders may be present in roots [28], although fungi associated with the FRE morphology appear to be phylogenetically distinct from putative ectomycorrhizal and saprotrophic Endogonales [28].

Comparative genome analysis showed that the MAG’s genome had many features in common with G-AMF. Some of these genomic features, including the low abundance of PCWDE and the absence of thiamine biosynthesis genes, were also seen in the Endogonales genomes and therefore aren’t likely to be specific, hallmark features of an adaptation to a symbiotic lifestyle. However, other signatures the MAG shares with G-AMF, including a high abundance of calcium transporters and the absence of fatty acid synthases, may point to physiological convergence in the way in which the symbioses operate. There were some notable differences between the MAG and G-AMF, with the MAG possessing a high abundance of chitinases, a complete vitamin B6 biosynthesis pathway, and low abundances of nitrate, ammonium, and copper transporters. These features may point to differences between G-AMF and M-AMF in the way they interact with the host plant and the wider environment.

It has been suggested that the transition from saprotrophic to mycorrhizal lifestyles involved loss of PCWDE, enabling fungi to exist within roots without digesting them [25]. The G-AMF symbiosis is estimated to have evolved 450 mya, and Glomeromycotan genomes possess substantially lower numbers of these genes than the fungi forming the more recently evolved ectomycorrhizal, ericoid mycorrhizal, and orchid mycorrhizal symbioses. Furthermore, G-AMF show signatures of dependence on host metabolism that are absent from more recently evolved symbioses, including fatty acid auxotrophy and an inability to synthesise thiamine and metabolise vitamin B6 [25]. These represent the ‘Missing Glomeromycota Core Genes’ (MGCG) [15–17].

Although genomic evidence indicates that G-AMF and M-AMF have independent origins [17], these symbioses co-occurred in the earliest land plants [7], and they continue to co-occur in roots of extant plants [22]. Additionally, while G-AMF and M-AMF exhibit some niche differentiation, evidence of overlapping ecological presence and similar responses to overarching edaphic and environmental variables have been noted [27, 28]. Furthermore, ultrastructural evidence points to similarities in the way in which G-AMF and M-AMF function at a cellular level [24]. Therefore, similar evolutionary pressures may have acted on these symbioses, which could have resulted in convergence of genomic traits. Indeed, many of the symbiotic genomic features shown by G-AMF were also displayed by the MAG. This included a reduced repertoire of PCWDE and the absence of several MGCG, including those related to thiamine and fatty acid biosynthesis.

Our analysis confirms the findings of Rosling et al. (2024) that Endogonales genomes have a low genomic PCWDE count and an incomplete thiamine biosynthesis pathway. Similarly, our MAG, which represents an early-diverging lineage within the Mucoromycotina and is possibly related to the Densosporales, also displays these features. This suggests that these traits may be present in multiple lineages within the Mucoromycotina and may not be exclusive adaptations to biotrophy. However, the Endogonales genomes possess fatty acid biosynthesis genes [17], whereas both the G-AMF genomes and the MAG lack these genes. This absence in both G-AMF and the MAG could indicate a convergent adaptation to biotrophy, potentially suggesting that host assimilate is provided as fatty acids in both symbioses.

The MAG had a number of features which distinguished it from G-AMF genomes. In particular, while there was a shared low abundance of CAZymes, the MAG possessed a greater repertoire of enzymes for degradation of organic substrates than typically seen in G-AMF genomes. This included the full complement of enzymes required for degradation of cellulose, which was absent in all G-AMF genomes. In notable contrast to G-AMF, the MAG had a high count of both chitin and mannan fungal cell wall degrading enzymes. The high chitinase count doesn’t appear to be specific to the M-AMF, with the putative ectomycorrhizal and saprotrophic Endogonales genomes showing a similar pattern, attributed to elevated numbers of CE4 and GH18, as seen previously in Rosling et al. (2024). However, the full complement of chitinases was present in the G-AMF genomes as well as in the MAG, so this feature may not have any functional significance. Relative to other mycorrhizal fungi (except ericoid mycorrhizal fungi), the MAG had a notably high count of genes for mannan degradation. These were absent or present in low numbers in most G-AMF genomes and, significantly, were also found in low abundance in the Endogonales genomes, therefore high abundance of mannan degradation genes may represent a distinguishing feature of FRE genomes. However, even within the G-AMF there appears to be some variation, with the Diversisporales genomes containing a higher number of mannan and chitin degrading enzymes than the Glomerales genomes, indicating a substantial range of genomic repertoires.

It is important to understand whether these contrasting enzymatic profiles contribute to niche partitioning between G-AMF and M-AMF in nutrient mobilisation from contrasting mineral and organic substrates. For example, high abundance of chitin and mannan degrading genes could indicate a role for M-AMF in the degradation of fungal necromass, while a complete cellulose degradation pathway and high abundance of cellulose and hemicellulose degrading enzymes could suggest involvement in saprotrophic breakdown of plant-derived materials, similar to some ECM fungi [81, 5]. Alternatively, it could reflect different strategies for enzymatic degradation of plant cell walls as they colonise and grow through host plant tissues and penetrate root cells.

Currently, there is no indication that soil organic matter has any influence on the distribution of G-AMF or M-AMF. In contrast to G-AMF, evidence indicates that M-AMF are more abundant in agricultural than natural habitats, indicating a preference for low organic matter soils [27]. However, these functional differences could support niche partitioning within these habitats.

A further distinguishing feature between the MAG and the G-AMF genomes is the presence of a full B6 biosynthesis pathway in the MAG, including pyridoxine kinase, which was missing in all the G-AMF genomes, as demonstrated in earlier studies [15, 16]. B6 is a cofactor required for amino acid biosynthesis, and it also plays a role as an antioxidant. Reactive oxygen species (ROS) scavenging systems such as B6 may be required by AMF to overcome host defences, allowing intracellular colonisation and the development of arbuscules [82, 83]. Components of the B6 biosynthesis pathway that G-AMF do possess are expressed throughout the fungal life cycle, but they are at the highest expression levels within the root [84]. Similarly, B6 plays a role in supporting the infection of root tissues by pathogens [85]. Differences in capacity for independent production of B6 between the MAG and G-AMF could point to differences in the way the fungi scavenge ROS, and potentially also differences in the way root colonisation and arbuscule development are regulated by the plant. There could also be implications for the specific mechanisms underlying M-AMF and G-AMF responses to external stressors which generate ROS such as temperature, osmotic stress, and nutrient limitation, and the independence of these responses to the plant’s own response [86].

Key physiological traits associated with G-AMF symbioses are the uptake of P from the soil, its translocation through hyphae to the root, and its transfer to the plant in exchange for carbon. Ultrastructural [24] and experimental evidence [18] suggest a similar role for M-AMF. A number of P transporters have been identified in G-AMF involved in P assimilation from soil, storage of P, and exchange of P with the host [87–89]. G-AMF genomes generally showed higher mean numbers of P transporters than ECM, while numbers in the MAG were low relative to other mycorrhizal genomes studied. While the significance of transporters to the functioning of symbionts will depend on a range of factors including their efficiency, our data doesn’t provide genomic support for particular specialisation in P nutrition by M-AMF.

Evidence also points to a potential role for G-AMF in N nutrition [90]. Indeed, G-AMF genomes contained higher numbers of nitrate transporters than the other mycorrhizal symbionts, while also sitting towards the higher end of both ammonium and glycine transporters within the genomes we analysed. However, further evidence of inter-group differentiation can be seen across the Diversisporales and Glomerales genomes, with the former containing especially high numbers of ammonium and glycine transporters. The MAG contained a lower number of all three categories of N transporters than all G-AMF genomes. While the MAG contained a typical number of ammonium transporters compared to the other mycorrhizal lifestyles, it contained a low number of nitrate and glycine transporters. Hoystead et al. (2023) suggested that M-AMF and G-AMF have complementary roles in host nutrition, with M-AMF predominantly involved in supply of N to the host and G-AMF in supplying P. Furthermore, Howard et al. (2024) showed a putative M-AMF displaying both a preference for, and increased transfer of, glycine and ammonium derived nitrogen sources. While there is evidence for the ability to transport varying forms of N, our MAG does not provide exceptional evidence for this relative specialism.

The very low abundance of Cu transporters in the MAG - the lowest across the entire dataset - is particularly striking. The three ectomycorrhizal Endogonales genomes also contain low numbers of Cu transporters, however the saprotrophic Endogonales sit near the mean. G-AMF may promote Cu uptake by the host plant [91] and arbuscules have high demands for Cu [92]. Furthermore, Cu transporters in G-AMF show differential expression profiles across intra- and extra-radical mycelium [92]. The extent to which M-AMF are involved in the supply of micronutrients to the host remains to be determined, but the evidence presented here could indicate that in contrast to G-AMF, this may not be an important function for M-AMF.

The high number of calcium transporters in both the MAG and G-AMF is a clear point of similarity between them, and of differentiation to all other groups. AMF have been implicated in transport of Ca to host plants [93], and our data could therefore indicate a role for both G-AMF and M-AMF in this function. However, Ca may also play a key role in the arbuscule lifecycle of both FRE and G-AMF. A clear link between a dramatic increase in calcium accumulation and arbuscule collapse has been demonstrated within G-AMF [94, 95]. A similar trend within FRE has also been documented [24, 96]. While it is unknown whether the Ca accumulated originates from the host plant or the fungi, the high number of Ca transporters within both G-AMF and the MAG may indicate that Ca accumulation is driven by the fungal symbionts, perhaps depositing Ca to cease the function of the arbuscule once it reaches maturity.

We must exercise some caution when interpreting these findings since we have constructed a relatively fragmented genome from a metagenome. However, our predicted level of contamination is only 1.5%, this suggests that we are unlikely to be assigning functions to the MAG that are not really present. In contrast, completion is potentially low at 83.7% so it is a definite possibility that we have missed a significant number of genes but most of the analysis above is at the level of gene families, if the missing genes are randomly distributed through these families we are unlikely to predict the absence of an entire family that is actually possessed by our MAG.

## Conclusion

In this study, we assembled and characterized a MAG of a putative fine root endophyte forming Mucoromycotina AMF from samples collected in Australia. Comparative analyses revealed key genomic features shared with related fungal taxa, shedding light on potential functional adaptations to the root-associated lifestyle. Our findings contribute to the growing understanding of Mucoromycotan-AMF diversity and their ecological roles, providing a valuable genomic resource for future research on endophytic symbioses and nutrient exchange in plant-fungal interactions. For both G-AMF and M-AMF, comprehensive analysis of genomic functional gene repertoires across the broad range of taxa found within roots is key to understanding the way that AMF communities support plant health, and the potential for engineering communities to promote beneficial traits. With the development of third generation long read sequencing technologies, genome assembly from metagenomes represents a promising route for understanding functions and traits associated with dark taxa in complex microbial communities, especially when combined with transcriptome sequencing [40] to understand the factors which regulate gene expression within plant tissues and the soil.

## Declarations

### Ethics approval and consent to participate

Not applicable

### Consent for publication

Consent for use of Figure 1d obtained from Suzanne Orchard. Credit given in the figure legend.

### Availability of data and materials

Raw 18S rRNA Miseq sequences are available at NCBI (BioProject ID PRJNA1179892), and Nanopore sequences at ENA (Experiment Accession ERX13330708). The FRE4583 sequence is available at NCBI (SUB14894546). The 18S rRNA phylogeny sequence alignment FASTA file and Newick tree file are available on PlutoF Biodiversity Platform (https://dx.doi.org/10.15156/BIO/3301225). The sequence of the MAG has been deposited on the Zenodo repository (https://doi.org/10.5281/zenodo.14144746).

### Authors’ contributions

GB, MHR, and CQ conceived the work and with RM obtained funding. MHR and PAD conducted the M-AMF enrichment work and the assessment of M-AMF colonisation of plant roots. SH, SR, and GB conducted sequencing and analysis of 18S rRNA genes. SR and CQ assembled the MAG. RHN, GM, and JC conducted the 18S rRNA gene phylogenetic analysis. JC, GM, SR, and GB performed the comparative genomic analyses. JC and GB wrote the manuscript. All authors edited the manuscript.

## Acknowledgements

We thank the Natural Environment Research Council (Grant NE/S010270/1), the Biotechnology and Biological Sciences Research Council (BBSRC) Midlands Integrative Biosciences Training Partnership (Grant BB/T00746X/1) and the Australian Research Council (Discovery Project DP180103157) for funding.

