## Supplementary material for "Comparative genomic analysis of a metagenome-assembled genome reveals distinctive symbiotic traits in a Mucoromycotina fine root endophyte arbuscular mycorrhizal fungus": Supp_Figures

### Slide 1
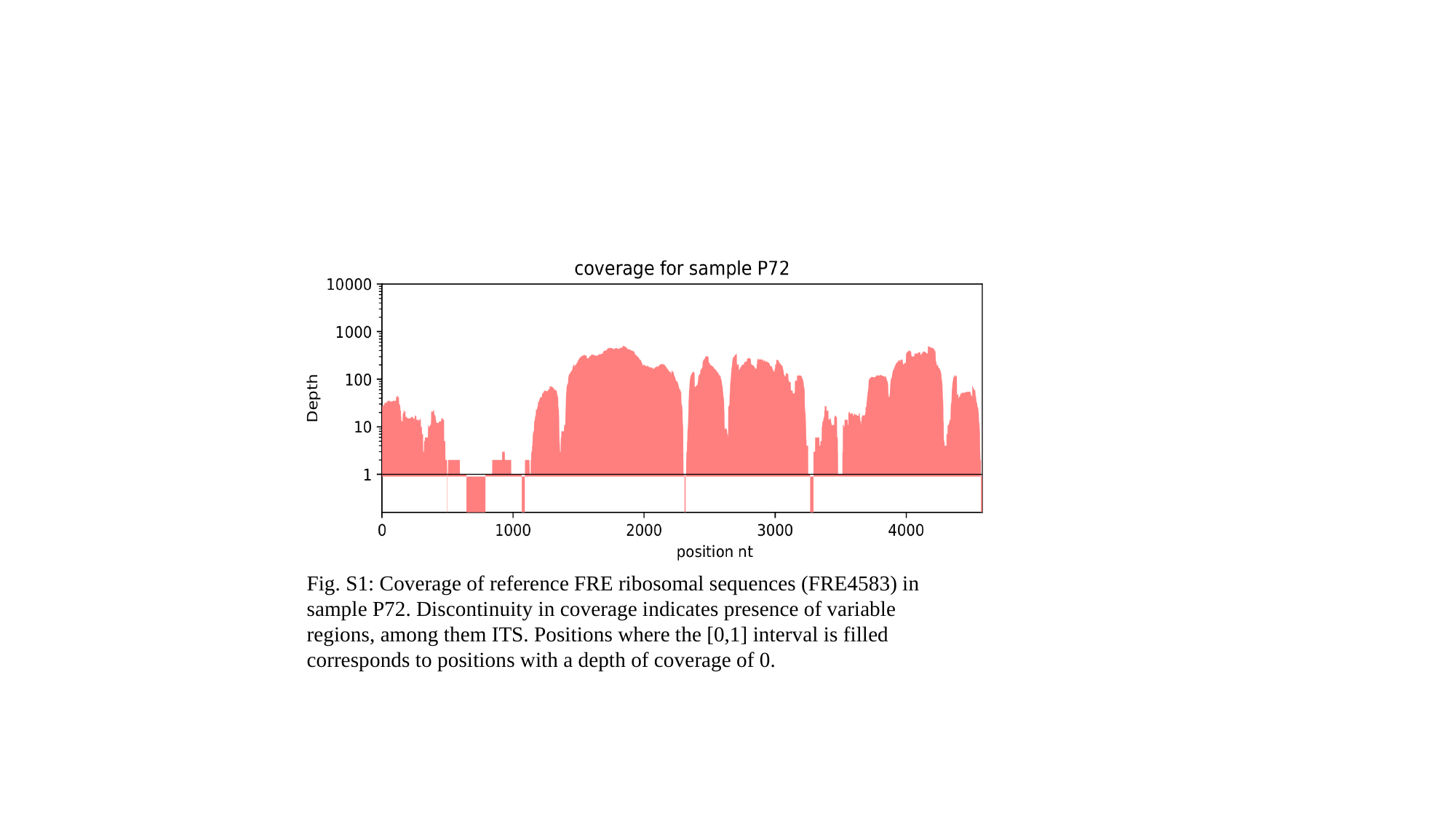

Fig. S1: Coverage of reference FRE ribosomal sequences (FRE4583) in sample P72. Discontinuity in coverage indicates presence of variable regions, among them ITS. Positions where the [0,1] interval is filled corresponds to positions with a depth of coverage of 0.

### Slide 2
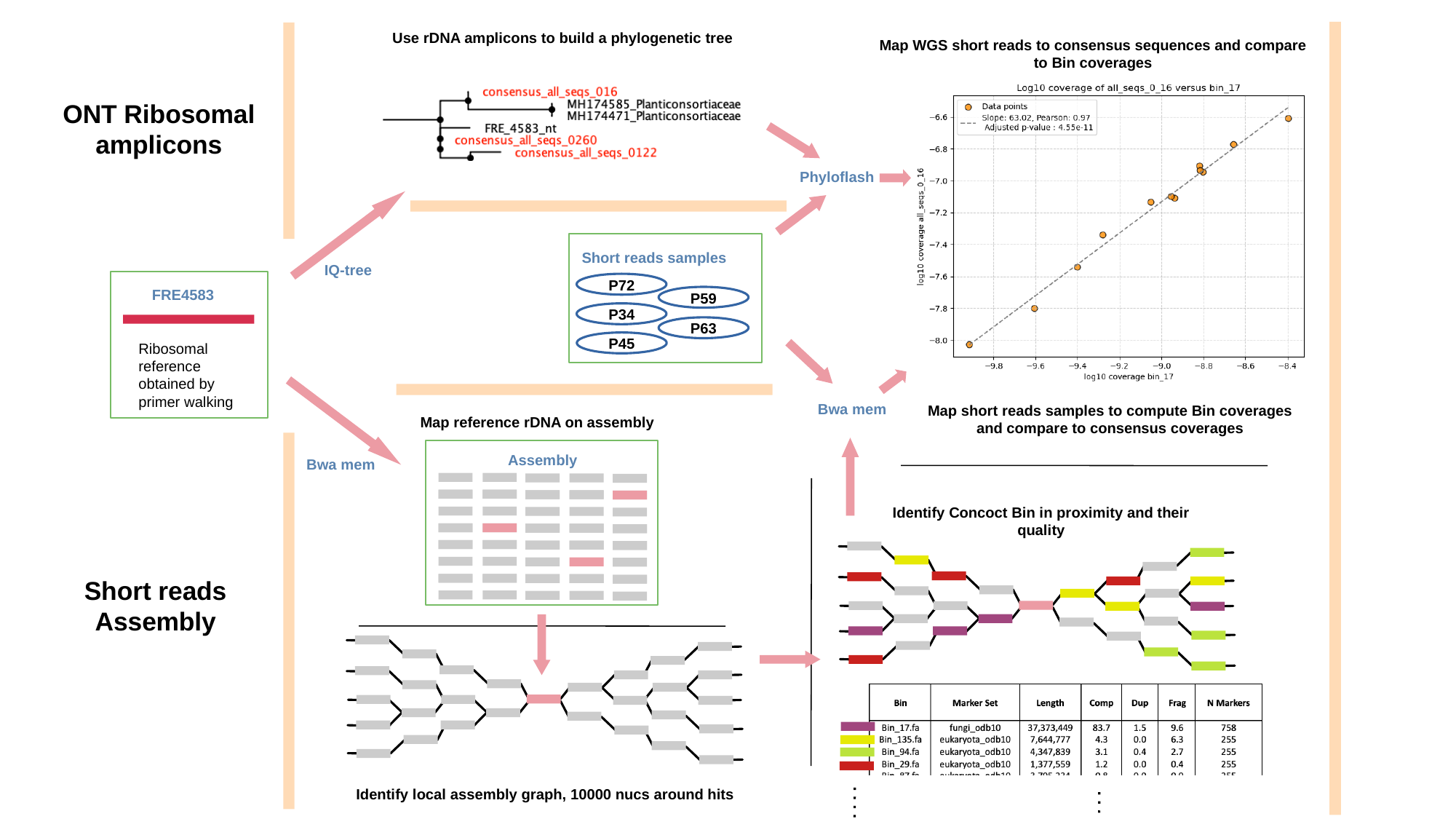

Use rDNA amplicons to build a phylogenetic tree
Map WGS short reads to consensus sequences and compare to Bin coverages
ONT Ribosomal
amplicons
Phyloflash
Short reads samples
IQ-tree
P72
FRE4583
P59
P34
P63
Ribosomal reference obtained by primer walking
P45
Bwa mem
Map short reads samples to compute Bin coverages and compare to consensus coverages
Map reference rDNA on assembly
Assembly
Bwa mem
Identify Concoct Bin in proximity and their quality
Short reads Assembly
Identify local assembly graph, 10000 nucs around hits
Fig. S2: Bioinformatic workflow leading to identification of the FRE MAG: FRE4583 is a reference sequence obtained by primer walking, it is used, from one hand to identify similar unitig in the assembly graph and on the other, to identify a clade of de novo ont ribosomal amplicon. As a part of the treatment of the assembly, the contigs are binned using CONCOCT and we focus on the few which are found to be in proximity of the flagged unitig.  The coverage of Bin_17, the only bin of good quality, is then compared to the coverage of the multiple ont amplicon from the FRE clade.  Consensus_all_seqs_016 is perfectly correlated with Bin_17, validating our finding.

### Slide 3
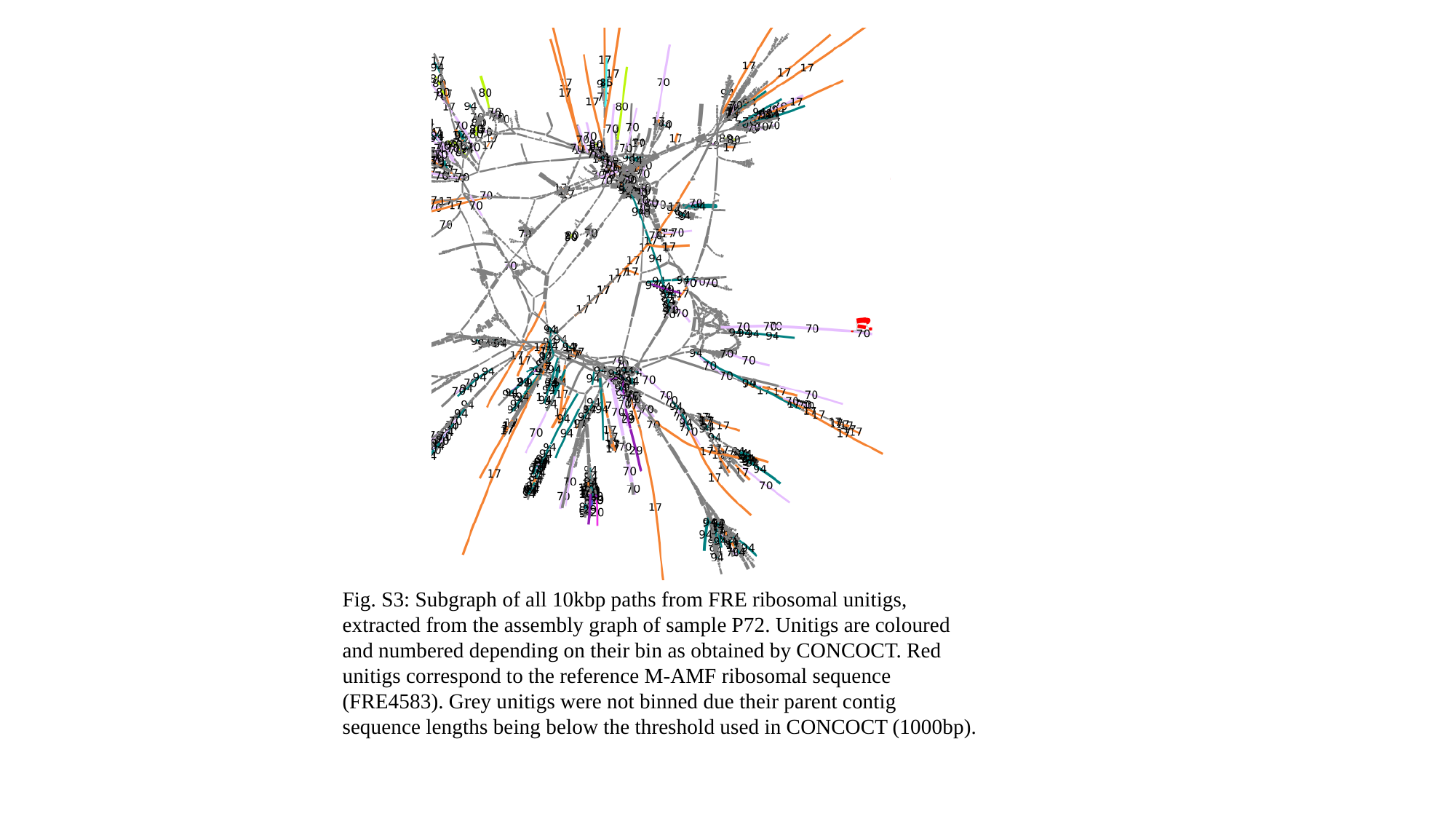

Fig. S3: Subgraph of all 10kbp paths from FRE ribosomal unitigs, extracted from the assembly graph of sample P72. Unitigs are coloured and numbered depending on their bin as obtained by CONCOCT. Red unitigs correspond to the reference M-AMF ribosomal sequence (FRE4583). Grey unitigs were not binned due their parent contig sequence lengths being below the threshold used in CONCOCT (1000bp).

### Slide 4
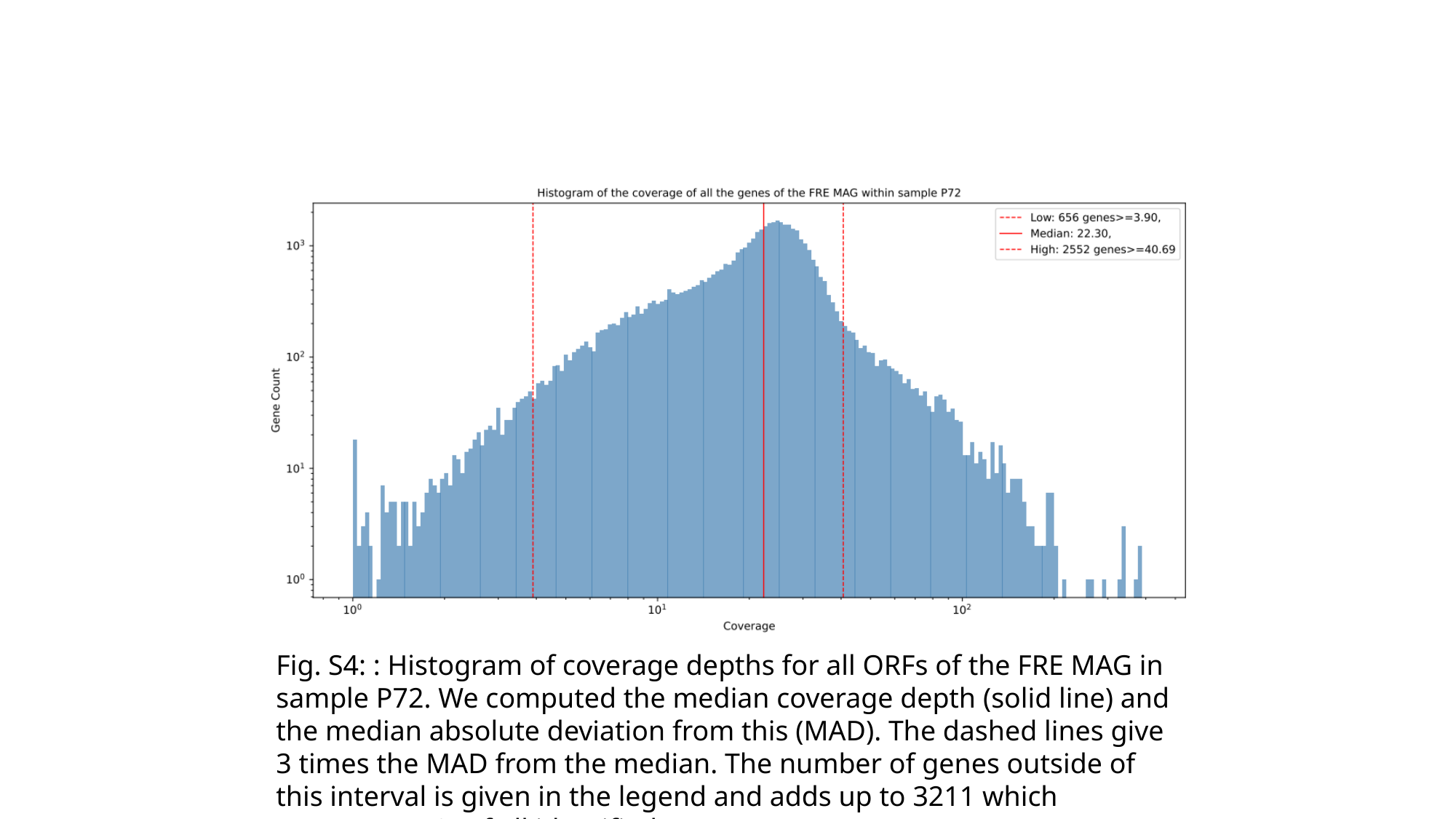

Fig. S4: : Histogram of coverage depths for all ORFs of the FRE MAG in sample P72. We computed the median coverage depth (solid line) and the median absolute deviation from this (MAD). The dashed lines give 3 times the MAD from the median. The number of genes outside of this interval is given in the legend and adds up to 3211 which represents 7% of all identified genes.

### Slide 5
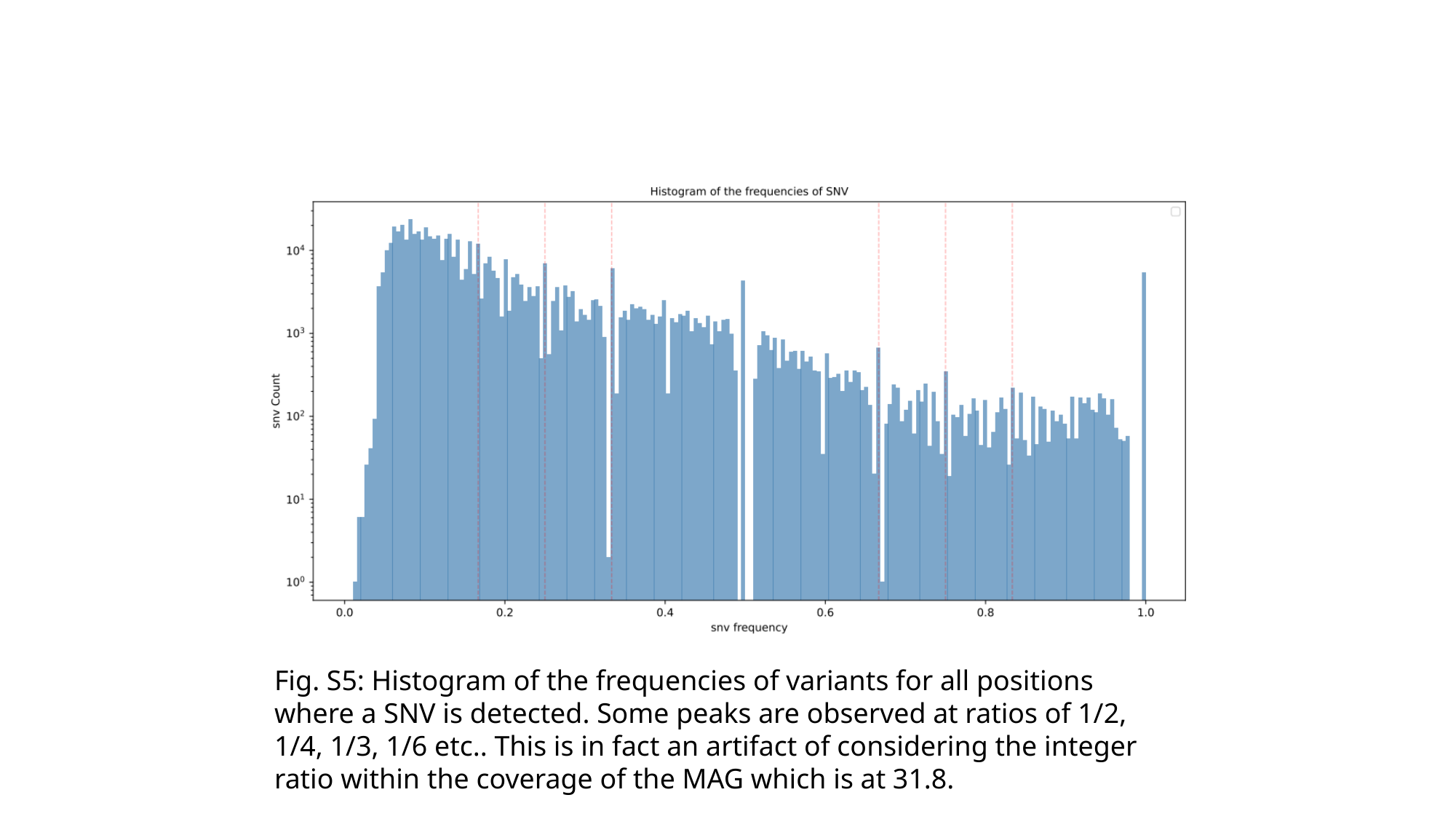

Fig. S5: Histogram of the frequencies of variants for all positions where a SNV is detected. Some peaks are observed at ratios of 1/2, 1/4, 1/3, 1/6 etc.. This is in fact an artifact of considering the integer ratio within the coverage of the MAG which is at 31.8.

### Slide 6
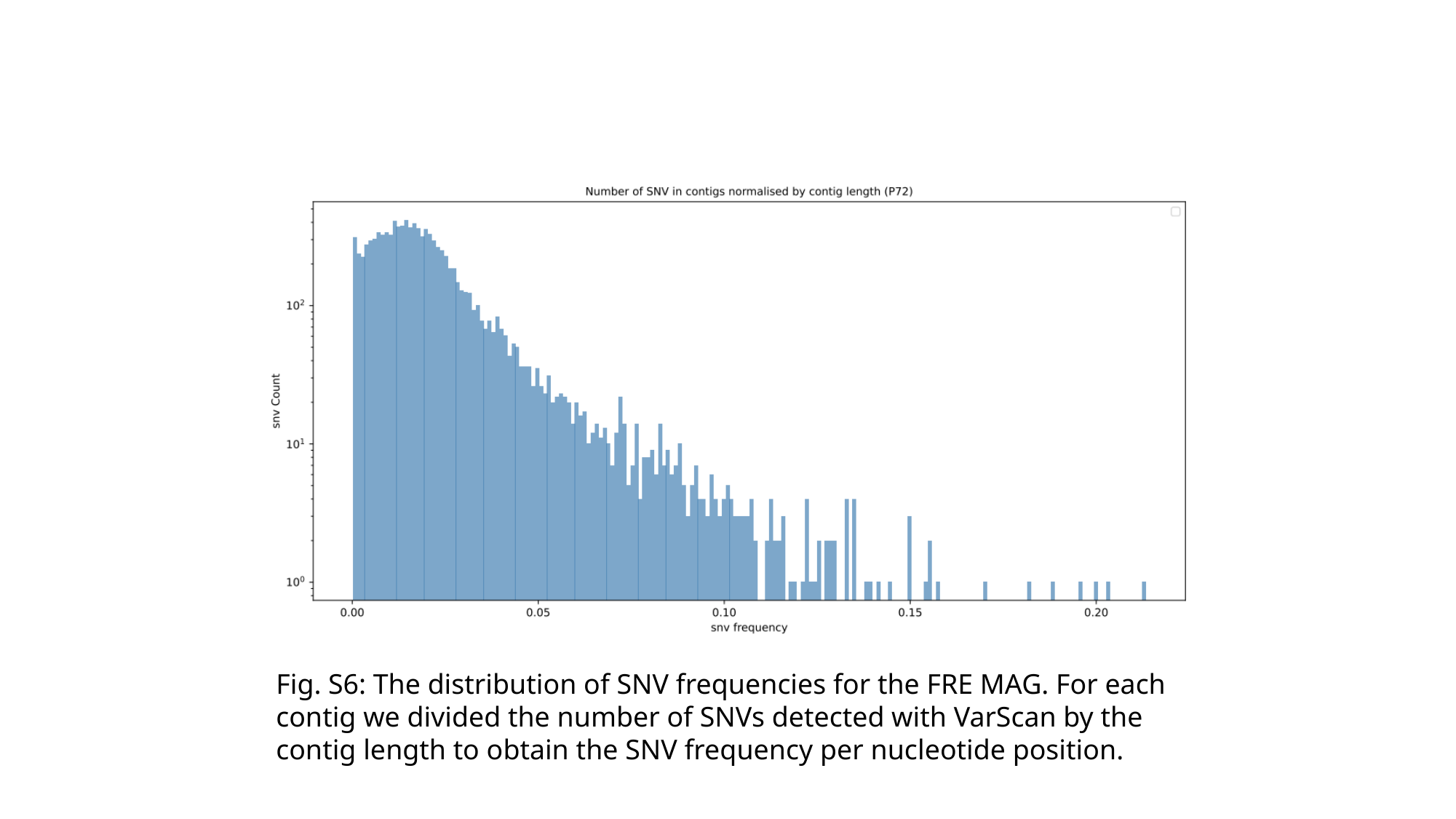

Fig. S6: The distribution of SNV frequencies for the FRE MAG. For each contig we divided the number of SNVs detected with VarScan by the contig length to obtain the SNV frequency per nucleotide position.

### Slide 7
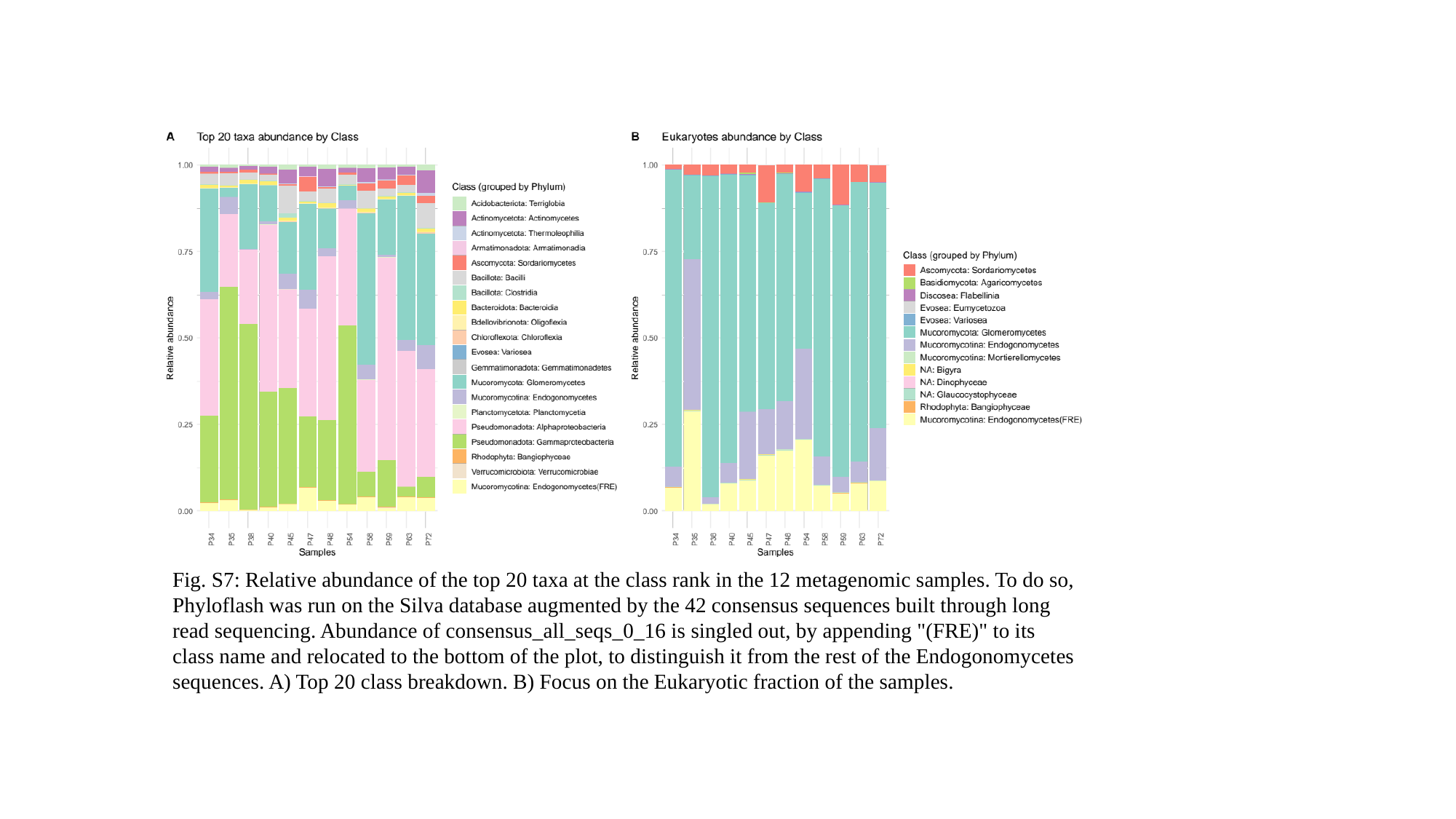

Fig. S7: Relative abundance of the top 20 taxa at the class rank in the 12 metagenomic samples. To do so, Phyloflash was run on the Silva database augmented by the 42 consensus sequences built through long read sequencing. Abundance of consensus_all_seqs_0_16 is singled out, by appending "(FRE)" to its class name and relocated to the bottom of the plot, to distinguish it from the rest of the Endogonomycetes sequences. A) Top 20 class breakdown. B) Focus on the Eukaryotic fraction of the samples.

### Slide 8
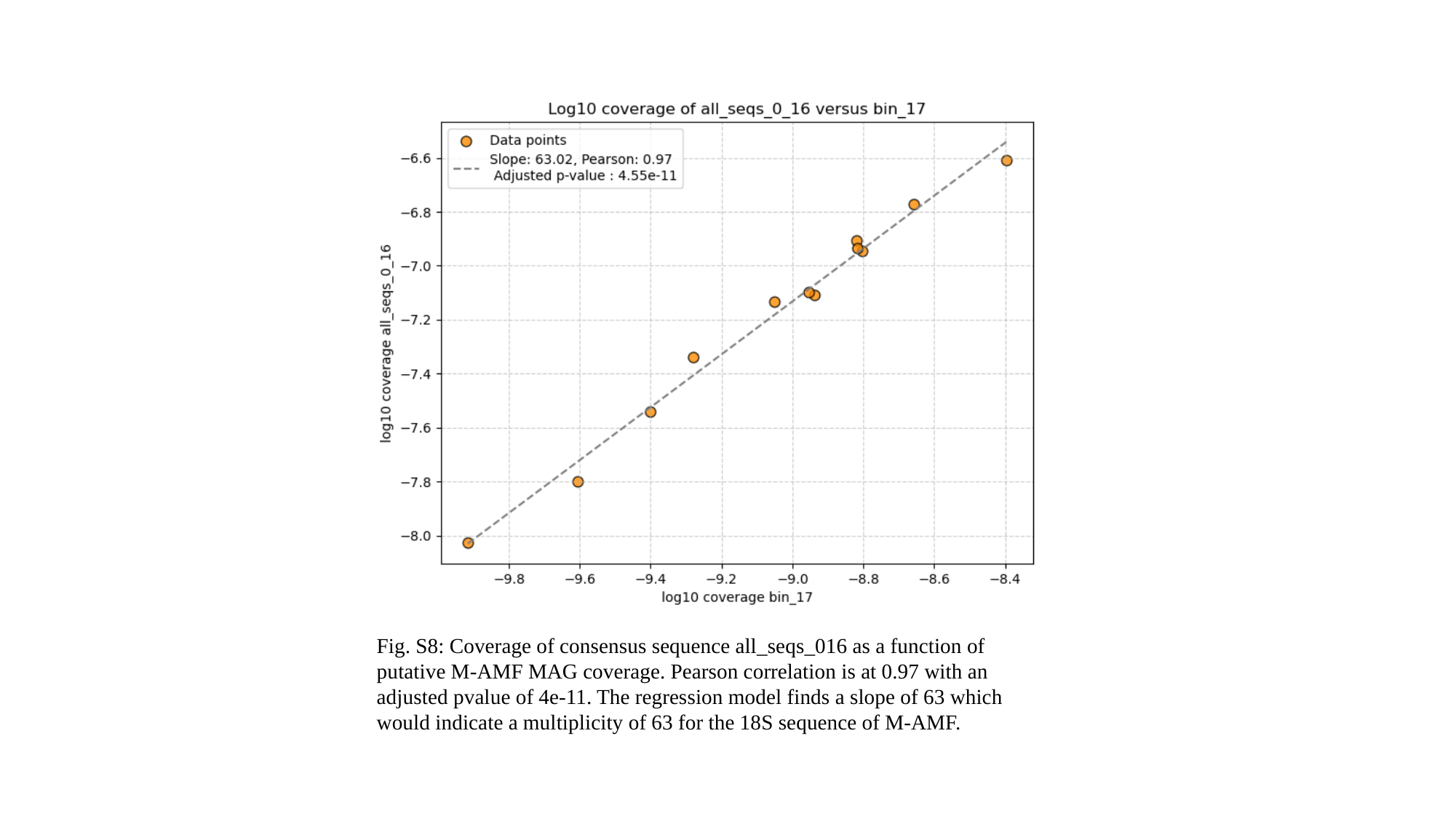

Fig. S8: Coverage of consensus sequence all_seqs_016 as a function of putative M-AMF MAG coverage. Pearson correlation is at 0.97 with an adjusted pvalue of 4e-11. The regression model finds a slope of 63 which would indicate a multiplicity of 63 for the 18S sequence of M-AMF.

### Slide 9
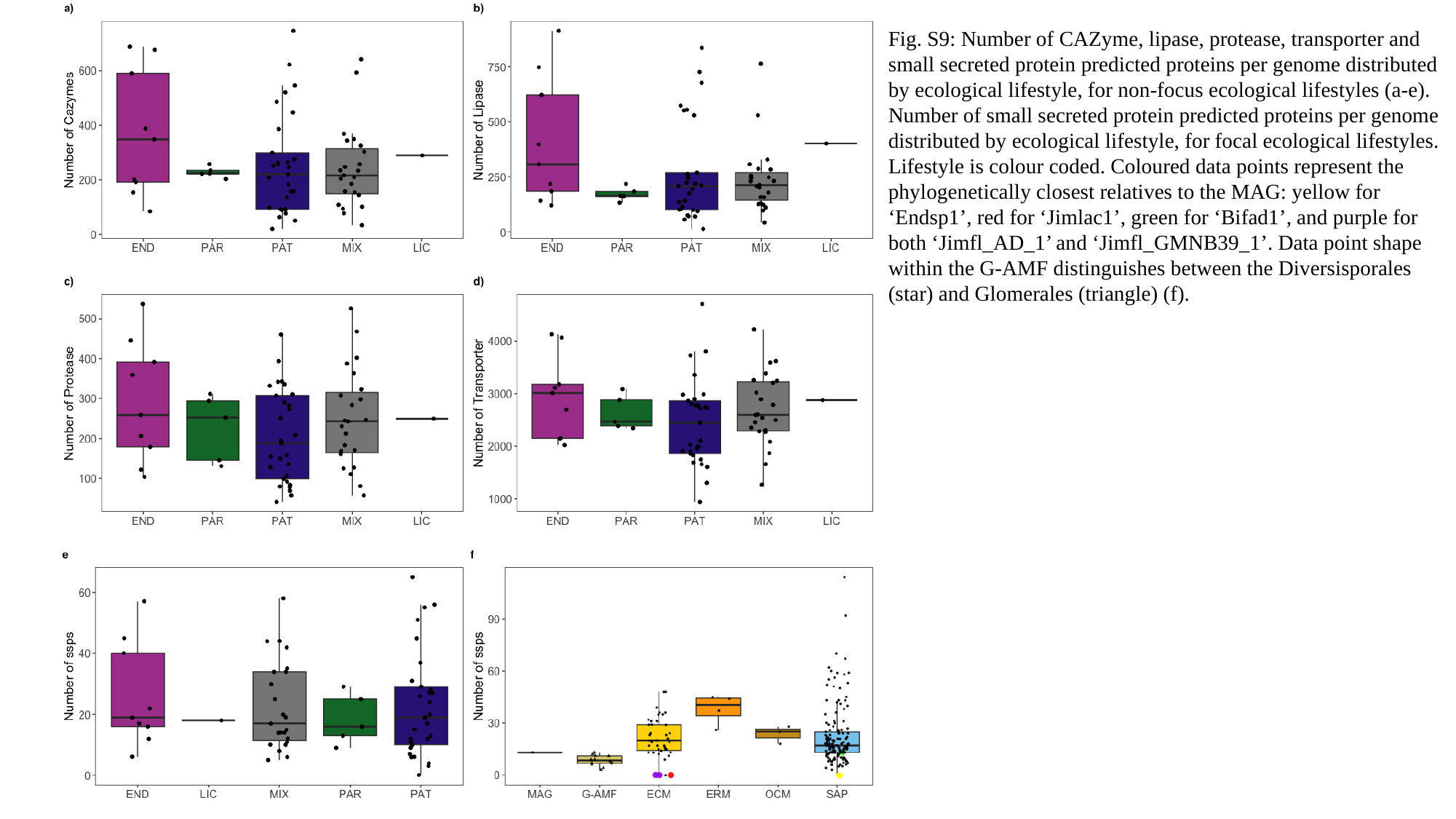

Fig. S9: Number of CAZyme, lipase, protease, transporter and small secreted protein predicted proteins per genome distributed by ecological lifestyle, for non-focus ecological lifestyles (a-e). Number of small secreted protein predicted proteins per genome distributed by ecological lifestyle, for focal ecological lifestyles. Lifestyle is colour coded. Coloured data points represent the phylogenetically closest relatives to the MAG: yellow for ‘Endsp1’, red for ‘Jimlac1’, green for ‘Bifad1’, and purple for both ‘Jimfl_AD_1’ and ‘Jimfl_GMNB39_1’. Data point shape within the G-AMF distinguishes between the Diversisporales (star) and Glomerales (triangle) (f).

### Slide 10
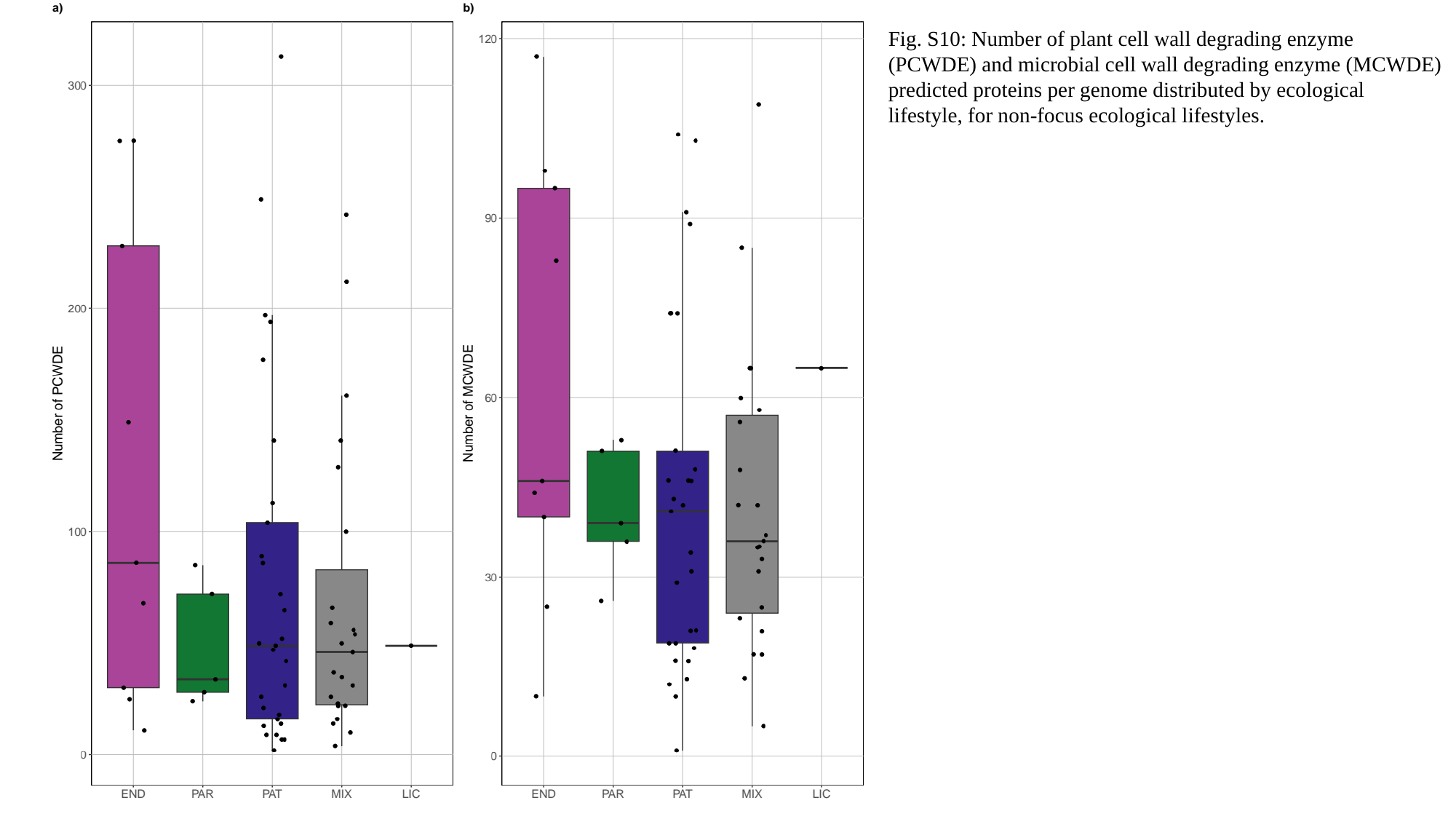

Fig. S10: Number of plant cell wall degrading enzyme (PCWDE) and microbial cell wall degrading enzyme (MCWDE) predicted proteins per genome distributed by ecological lifestyle, for non-focus ecological lifestyles.

### Slide 11
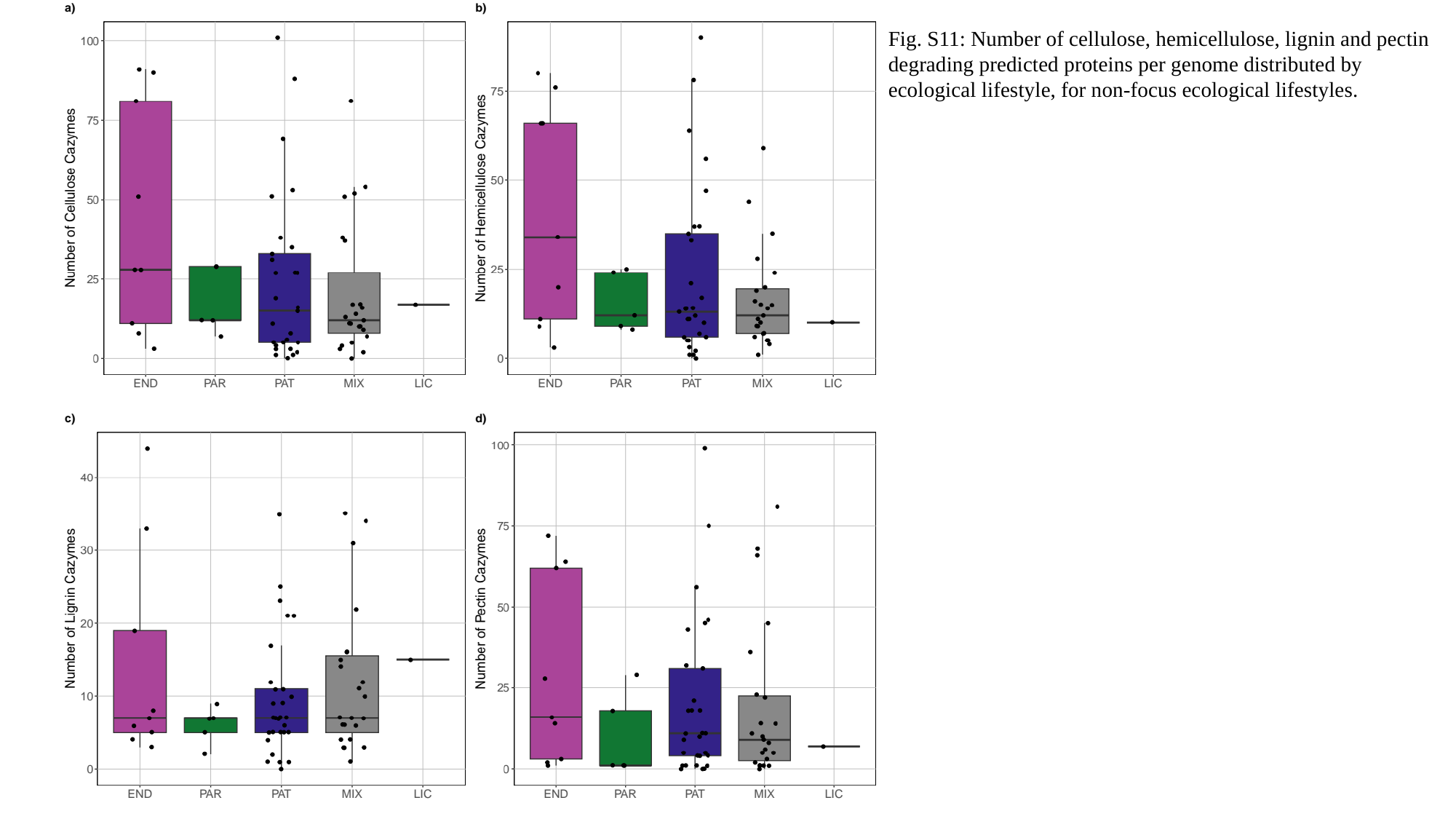

Fig. S11: Number of cellulose, hemicellulose, lignin and pectin degrading predicted proteins per genome distributed by ecological lifestyle, for non-focus ecological lifestyles.

### Slide 12
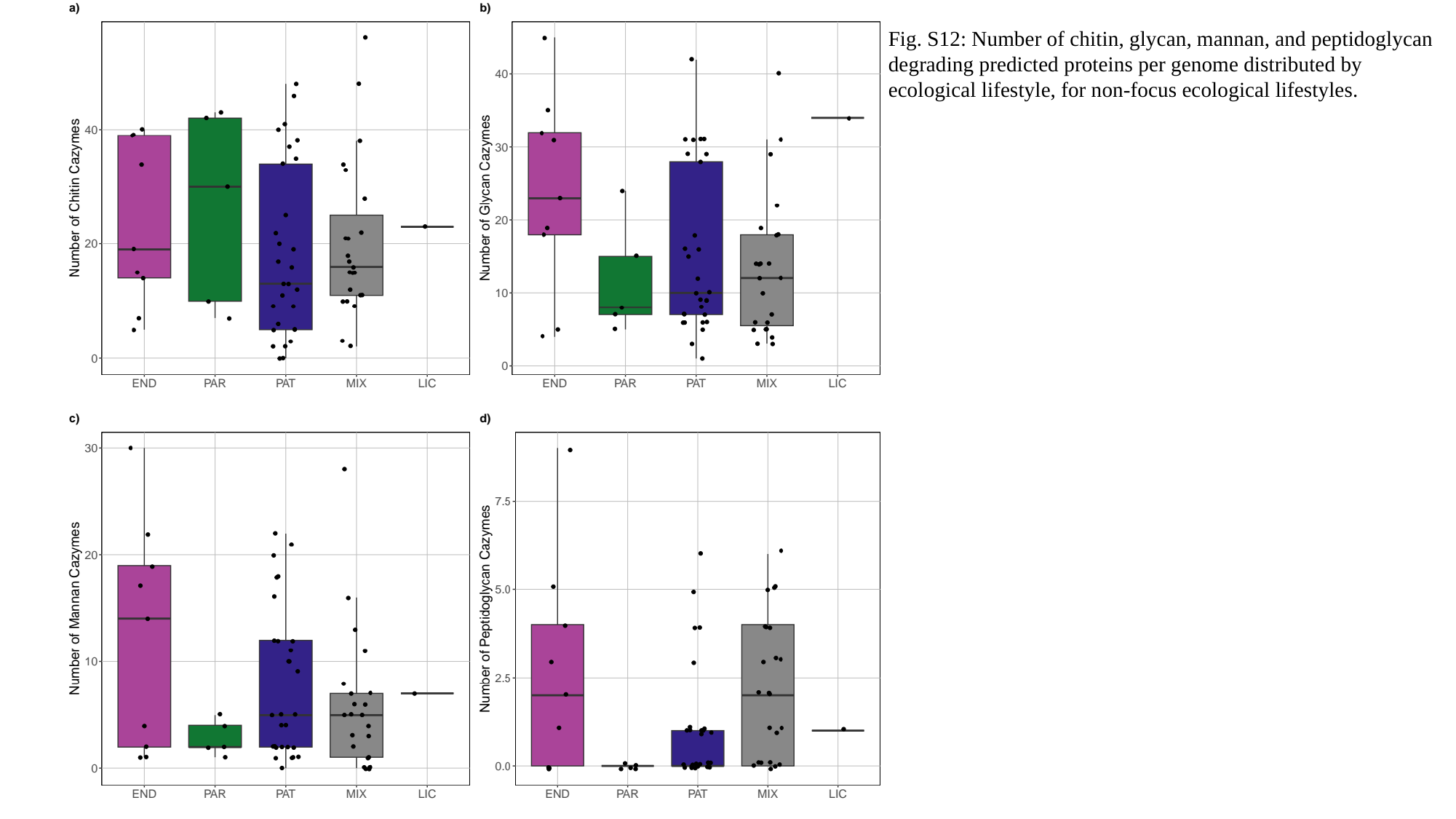

Fig. S12: Number of chitin, glycan, mannan, and peptidoglycan degrading predicted proteins per genome distributed by ecological lifestyle, for non-focus ecological lifestyles.

### Slide 13
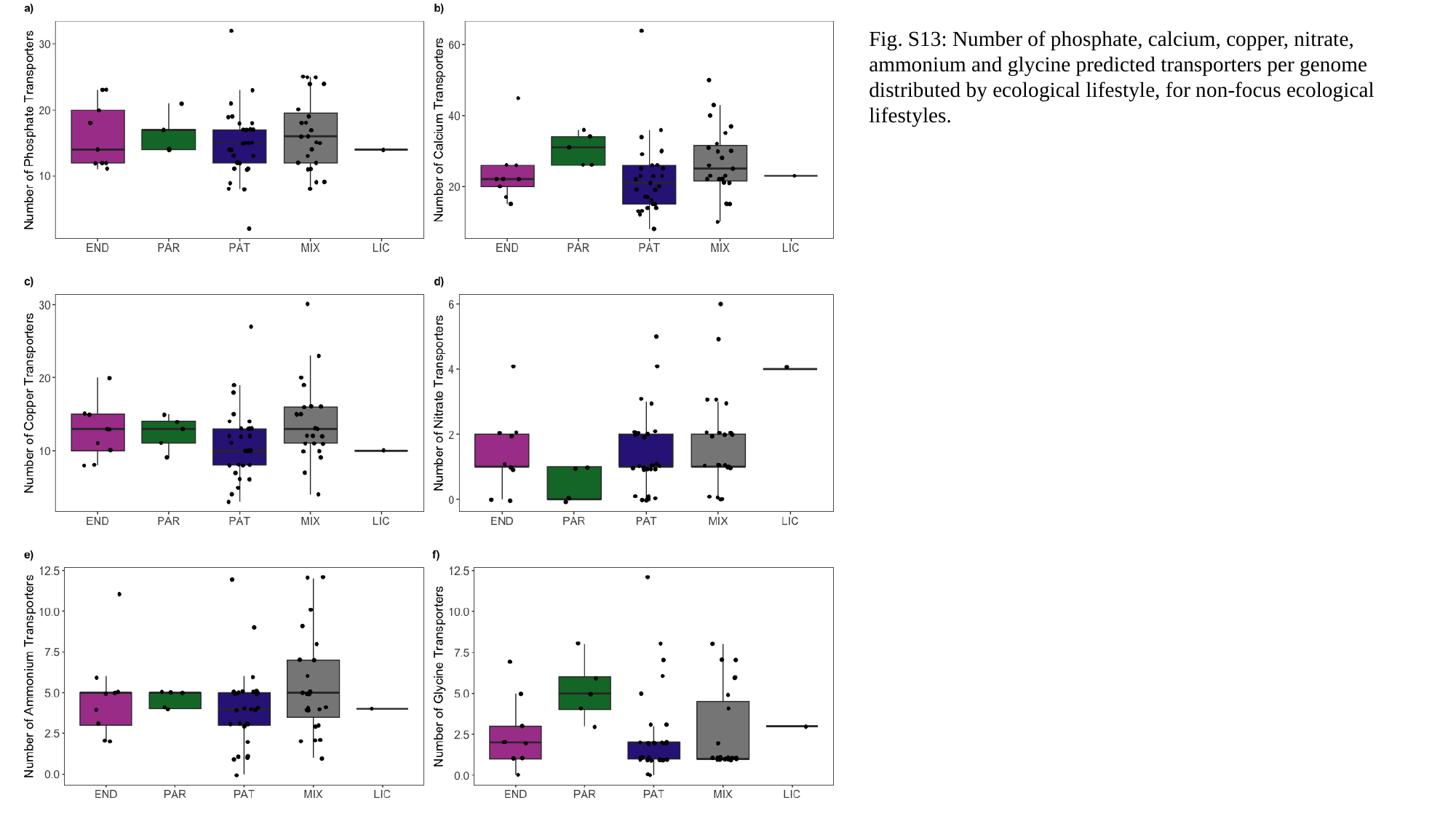

Fig. S13: Number of phosphate, calcium, copper, nitrate, ammonium and glycine predicted transporters per genome distributed by ecological lifestyle, for non-focus ecological lifestyles.
