## Supplementary material for "Comparative genomic analysis of a metagenome-assembled genome reveals distinctive symbiotic traits in a Mucoromycotina fine root endophyte arbuscular mycorrhizal fungus": Supp_Tables

### Slide 1
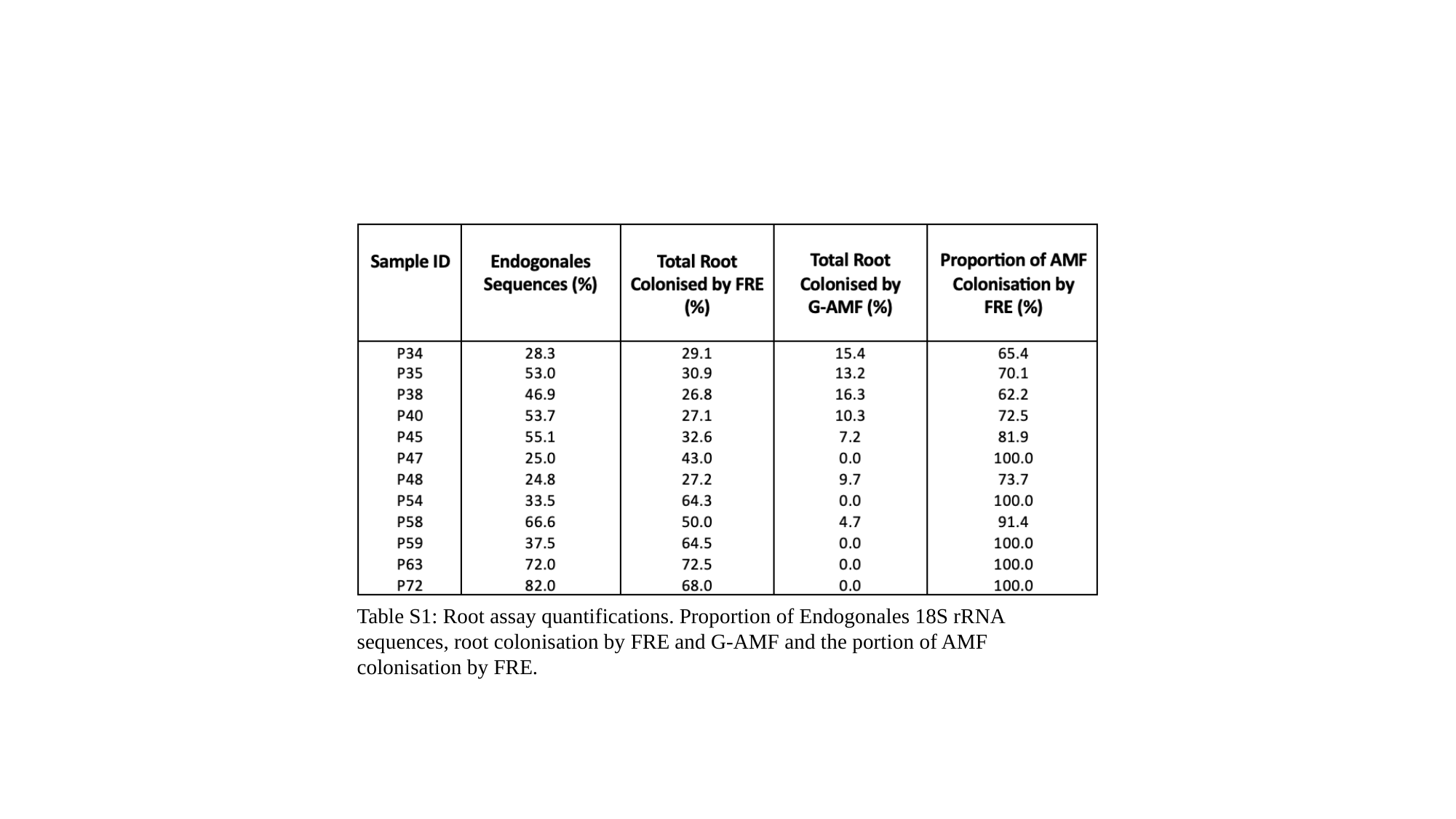

Table S1: Root assay quantifications. Proportion of Endogonales 18S rRNA sequences, root colonisation by FRE and G-AMF and the portion of AMF colonisation by FRE.

### Slide 2
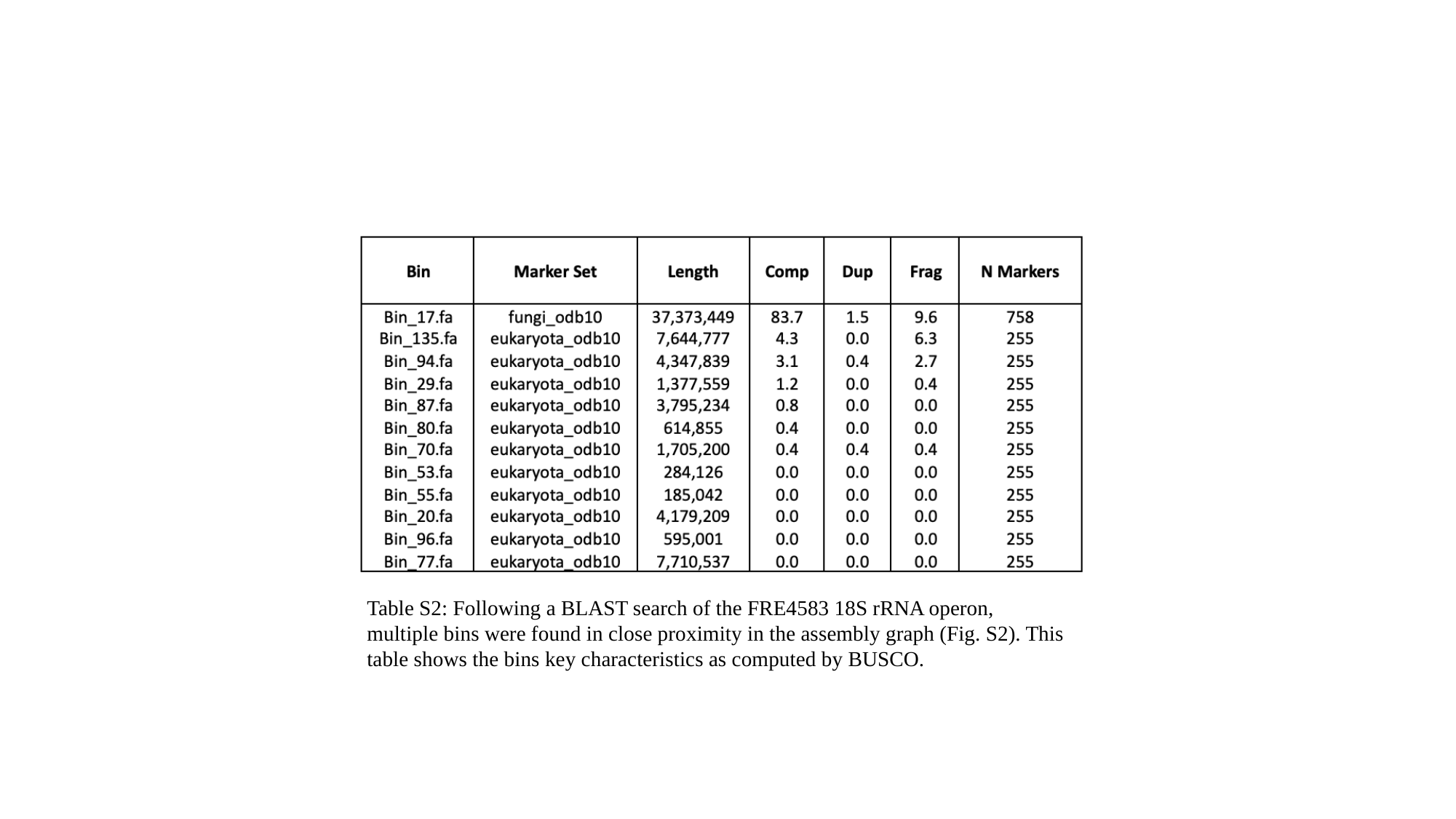

Table S2: Following a BLAST search of the FRE4583 18S rRNA operon, multiple bins were found in close proximity in the assembly graph (Fig. S2). This table shows the bins key characteristics as computed by BUSCO.

### Slide 3
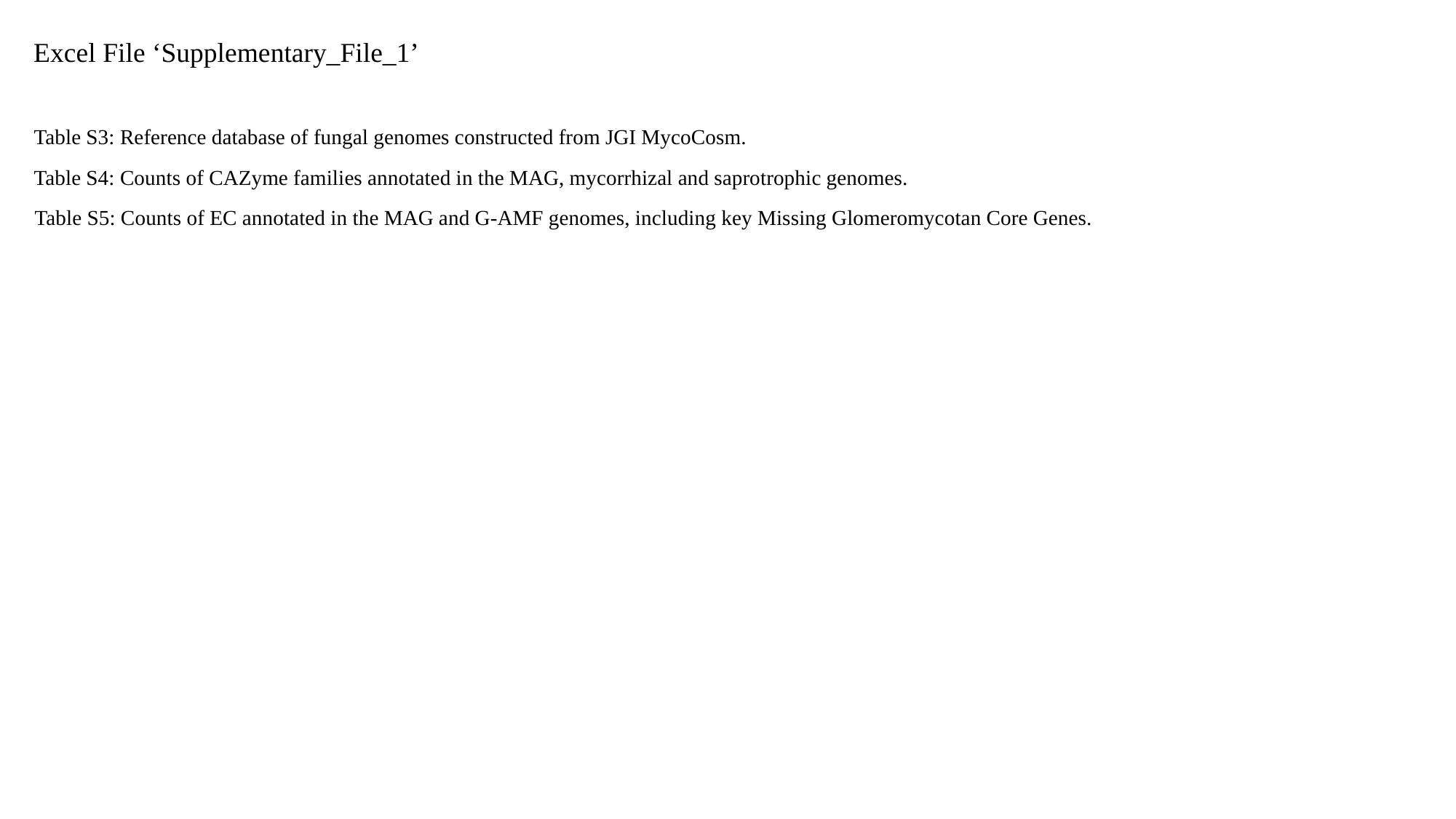

Excel File ‘Supplementary_File_1’
Table S3: Reference database of fungal genomes constructed from JGI MycoCosm.
Table S4: Counts of CAZyme families annotated in the MAG, mycorrhizal and saprotrophic genomes.
Table S5: Counts of EC annotated in the MAG and G-AMF genomes, including key Missing Glomeromycotan Core Genes.
