## Supplementary material for "Comparative genomic analysis of a metagenome-assembled genome reveals distinctive symbiotic traits in a Mucoromycotina fine root endophyte arbuscular mycorrhizal fungus": Supp_Text

**Supplementary Text**

**Supplementary Materials and Methods**

**Soil collection and sample preparation**

The field contained winter-active annual grasses and legumes and supported a low-intensity free-range beef cattle grazing enterprise. Soil collection targeted a sandy loam soil with a pH in CaCl2 of 5.4. Soil was wet-sieved through a stack of four sieves with pore sizes ranging from 2 mm down to 20 µm. The material collected on each sieve was retained and added to pasteurized, dried, washed coarse river sand at a ratio of either 1:27 or 1:81 w/w. Free-draining pots were filled with 1 kg of each treatment (n=4) and placed in a temperature-controlled glasshouse. Four plants of T. subterraneum L. with appropriate rhizobia added were grown in each pot, which were maintained at 80% field capacity by watering to weight and received phosphorus-free, but otherwise complete, nutrient solution once a week. After 8 weeks, plants were harvested, and roots gently washed clean. The four root systems from each pot were allocated to one of two future activities: staining for assessment of AMF colonization (stored in 70% ethanol) or DNA extraction (frozen at -20°C). Roots were cleared and stained using the ink-vinegar method of Vierheilig et al. (1998) and assessed using a modified version of the magnified intercept method described by McGonigle et al. (1990), where the root at each grid-line root intercept was recorded as either free of all colonization, colonized by G-AMF only, colonized by M-AMF only, or colonized by both M-AMF and G-AMF.

**Long read nanopore sequencing**

An initial PCR was performed with adapter-tagged oligos, and the PCR products were cleaned using Ampure SPRI beads. Barcodes were added using a second PCR with an initial denaturation at 95°C for 3 minutes followed by 30 cycles of denaturation at 98°C for 15 seconds, 60°C annealing for 15 seconds, and 65°C extension for 150 seconds, with a final extension at 65°C for 6 minutes. Following a clean-up with SPRI beads, the samples were used for sequencing on a SpotON Nanopore flow cell following the manufacturer’s instructions.

**Reference genome access and ecological lifestyle annotation**

We acknowledge that this was not an exhaustive list of fungal genomes accessible on JGI MycoCosm, in that there were genomes representing further species within Ascomycota and Basidiomycota that were not included in the analyses. Additionally, with our primary focus being the provision of comparable context for the MAG, we aimed to obtain a broadly representative database and not an all-encompassing list.

To achieve this, fungal species were coded as follows. First, ecological classifications were accessed from functional groups recorded on JGI MycoCosm (e.g., https://mycocosm.jgi.doe.gov/Ectomycorrhizal_fungi/Ectomycorrhizal_fungi.info.html). Secondly, for genomes included in previous comparative genomics analyses, the classifications reported in those studies were used ([3, 4, 17, 18]). Any remaining genomes without classifications were then manually curated, as informed by published evidence. Similar to previous studies, we allowed a fungal species to occupy multiple ecological lifestyles when there was evidence of this.

Of particular note were the 5 Mucoromycotina genomes within the dataset: Endogone sp. ‘Endsp1’ is a putative saprotroph found on fallen oak logs in Florida, USA [3]. Jimgerdemannia lactiflua, ‘Jimlac1’ is a putative ectomycorrhizal fungus isolated from under Pseudotsuga menziesii in Oregon, USA [3]. Two genomes of the putative ectomycorrhizal fungus Jimgerdemannia flammicorona: isolate ‘Jimfl_GMNB39_1’ from under Picea abies and Picea pungens in Michigan, USA [3], and isolate ‘Jimfl_AD_1’ from under Pinus strobus in Biella, Italy [3]. Bifiguratus adelaidae ‘Bifad1’, a soil-dwelling putative saprotroph isolated from a Pinus taeda plantation in North Carolina, USA [5].

**Annotation of MAG and reference genomes**

To identify CAZymes, fungal proteomes were queried against an HMM profile database built from CAZy sequences downloaded from the dbCAN2 database (https://bcb.unl.edu/dbCAN2/download/Databases/V11/dbCAN-HMMdb-V11.txt) [6] using HMMer v 3.3.2 [7]. Protein hits were subsequently filtered with an e-value threshold of 1e-15 and an alignment threshold of 50% of the domain query. To identify proteases, fungal proteomes were queried against the MEROPS peptide database (https://ftp.ebi.ac.uk/pub/databases/merops/current_release/merops_scan.lib) using BLASTp. Protein hits were subsequently filtered with an e-value threshold of 1e-10. To identify lipases, fungal proteomes were queried against the Lipase Engineering Database v4.1.0 using DIAMOND v2.1.8 [8] with the “--sensitive” option. Protein hits were subsequently filtered with an e-value threshold of 1e-5. To identify transporters, fungal proteomes were queried against the TCDB transporter database [9] using BLASTp. Protein hits were subsequently filtered with an e-value threshold of 1e-10. Secreted proteins were identified using a custom pipeline as reported previously [10]. Briefly, signal peptides were identified with SignalP v4.1 [11], extracellular localization was predicted with WoLFPSort v0.2 [12], transmembrane helices were identified with TMHMM v2.0 [13], secretory pathway association was inferred with TargetP v2.0 [14], and the absence of a KDEL motif in the C-terminal region was confirmed with PS-SCAN revision 1.86 [15]. Small-secreted proteins were identified as secreted proteins < 300 amino acids with no CAZyme, lipase, protease, or transporter annotation. Functional annotation of putative small-secreted proteins was achieved with PfamScan v1.6 [16], an e-value threshold of 1e-15, and an alignment threshold of 50% of the domain query. For each functional protein category identified, a single best hit per locus was retained by selecting the hit with the smallest e-value and greatest alignment of the domain query.

**Supplementary Results**

**CAZyme Profile**

G-AMF genomes contained significantly (p < 0.01) lower mean numbers of glycan-degrading genes (1) compared to the mean of the saprotroph (14) and other mycorrhizal lifestyles (15 to 42) and, in fact, were only detected in the two Diversisporales genomes (Fig. 7b). In contrast, the MAG contained 3 glycanases, consistent with a low number of glycanases in the Endogonales genomes (2 to 8). The MAG only contained 2 copies of GH17 and 1 copy of GH5_9 (Table S4). The only two other genomes to contain single copies of these were ‘Jimfl_AD_1’ and ‘Jimfl_GMNB39_1’. All the other Endogonales genomes contained 1 or 2 copies of both GH72 and GH81 (Table S4). The Diversisporales genomes only contained low counts of GH5_9 and GH81 (Table S4).

Low numbers of peptidoglycanases were found in the genomes of all lifestyles, with significantly larger mean numbers in ERM (8, p < 0.01) compared to the ECM (3) and saprotroph (2) genomes (Fig. 7d). The G-AMF genomes had a comparable mean number of peptidoglycanases (4) to the MAG (3), while the corresponding numbers in the Endogonales genomes were also low (0 to 3). The MAG only contained 2 copies of GH24 and a single copy of GH89 (Table S4). Similarly, the other Endogonales genomes were sparse in peptidoglycanases. Specifically, the ‘Jimfl_GMNB39_1’ genome contained no peptidoglycanases, while ‘Jimfl_AD_1’ and ‘Jimlac1’ only possessed a single copy of GH89 and CMB50, respectively. ‘Endsp1’ also contained 2 copies of CBM50 and a single copy of GH24. The G-AMF genomes also contained a small number of peptidoglycanases, mostly consisting of GH24 (Table S4).

The MAG contained lower numbers of PCWDE than the endophyte, parasite, pathogen, and mixed lifestyles (Fig. S10), but had higher numbers of MCWDE than the parasite, pathogen, and mixed lifestyles, with similar numbers to endophytes (Fig. S10).

Numbers of cellulose, hemicellulose, and pectin-degrading enzymes in the MAG were typically lower than the mean in endophyte genomes (Fig. S11), and numbers of lignin-degrading genes were similarly notably lower than the endophyte, parasite, pathogen, and mixed lifestyles (Fig. S11). Furthermore, the MAG contained notably higher than the mean number of chitinases found in endophytes, parasite, pathogen, and mixed lifestyles (Fig. S11).
